# Human Arm Redundancy - A New Approach for the Inverse Kinematics Problem

**DOI:** 10.1101/2023.02.20.528995

**Authors:** Avi Barliya, Nili Krausz, Hila Naaman, Enrico Chiovetto, Martin Giese, Tamar Flash

## Abstract

The inverse kinematics problem deals with the question of how the nervous system coordinates movement to resolve redundancy, such as in the case of arm reaching movements where more degrees of freedom are available at the joint versus hand level. In particular, this work focuses on determining which coordinate frames can best represent human movements, allowing the motor system to solve the inverse kinematics problem in the presence of kinematic redundancies. In particular, in this work we used a multi-dimensional sparse source separation method called FADA to derive sets of basis functions (here called sources) for both the task and joint spaces, with joint space being represented in terms of either the absolute or anatomical joint angles. We assessed the similarities between the joint and task sources in each of these joint representations. We found that the time-dependent profiles of the absolute reference frame’s sources show greater similarity to those of the corresponding sources in the task space. This result was found to be statistically significant. Hence, our analysis suggests that the nervous system represents multi-joint arm movements using a limited number of basis functions, to allow for simple transformations between task and joint spaces. Importantly, joint space seems to be represented in terms of an absolute reference frame to achieve successful performance and simplify inverse kinematics transformations in the face of the existing kinematic redundancies. Further studies will be needed to determine the generalizability of this finding and its implications for neural control of movement.

## Introduction

Human motor behavior is extremely rich, exhibiting a large variety of possible movements. However, the question of how the nervous system plans movement remains one of the most debated issues in the field of motor control. Human arm movements are of great interest given their particular versatility and adaptability, enabling the performance of numerous different motor tasks. To a large extent this versatility is due to the inherent kinematic redundancy of the human arm. Kinematically redundant limbs or manipulators are structures that have more degrees of freedom (DoFs) than those required for the performance of a specified task. Redundancy is highly beneficial when the need arises to overcome obstacles, adapt to environmental changes, or overcome fatigue. An excess number of DoFs can allow for motor tasks to be accomplished in many different ways.

While excess DoFs can prove useful, they raise the question of how the central nervous system (CNS) overcomes the computational problems associated with the neural control of redundant DoFs. Specifically, it is not fully understood how the CNS resolves kinematic redundancies to produce appropriate motor commands that can activate different limb muscles and produce forces and torques necessary to move the limb along a desired trajectory. This question was introduced by ***Bernstein (1967***) and was termed the “excess degrees of freedom problem”.

Redundancy, though, is not limited to the workspace or task levels. In general, the motor control hierarchy for planning movement has four levels: end-point (or task) variables (e.g. hand positions and velocities), joint angles and angular velocities, muscle activations, and neural activity patterns within cortical, sub-cortical, and spinal regions of the nervous system. It should be noted that redundancy appears at all of these levels. Firstly, most motor tasks can be performed using multiple possible end-effector postures and trajectories. Meanwhile, redundancy also exists at the level of joint rotations because the number of joint DoFs is larger than the number of end-effector DoFs. Moreover joint postures and rotations are controlled by generating appropriate joint torques produced by many alternative patterns of muscle activations. Due to the overlapping muscles’ actions and the ability to co-contract more muscles than are mechanically needed, the same arm configurations can be achieved through the generation of many different muscle activation patterns, which are determined at the neuronal level. Additionally, a rule of thumb is that a higher level in the motor hierarchy controls a greater number of state variables than the level below. Thus, at each level in the motor hierarchy (i.e. neural, muscular, joint, and task levels) these commands or states uniquely prescribe states at the level below. For example, joint positions prescribe the hand end-point position, and many different combinations of muscle commands can prescribe a particular joint position and limb configuration.

The inverse-kinematics (IK) problem is concerned with investigating how the CNS resolves this redundancy, for example by defining a unique arm configuration to achieve a desired hand location (in spite of the existing kinematic redundancy). Many researchers have tackled this problem and numerous approaches to finding the solution have been proposed. For example, ***Soechting and Terzuolo (1986***) addressed the inverse kinematics problem in the context of elliptical hand drawing movements. Particularly, they proposed a straightforward algorithm suggesting that elliptical hand trajectories result from oscillatory patterns of joint rotations, whereby the phase shifts between the different joint oscillations were assumed to be used to determine the geometrical form and orientation of the drawing plane. Others tried to apply methods from robotics involving the use of the generalized inverse of the manipulator or arm Jacobian (***Whitney, 1969***; ***Shamir, 1995***; ***Walker et al., 2008***; ***Siciliano, 1990***).

Cyclic drawing movements, however, pose some difficulty to these traditional approaches. For drawing a cyclic shape in which the end-effector returns to the initial point, these solutions produce non-repeatable trajectories in joint-space (***Klein and Huang, 1983***). Many additional studies have tried to solve this problem of joint repeatability by suggesting a variety of inverse kinematics algorithms that overcome this problem (***Hollerbach and Suh, 1987***; ***Baker and Wampler, 1987***; ***Shamir and Yomdin, 1988***; ***Klein and Kee, 1989***; ***Mussa-Ivaldi and Hogan, 1991***; ***Roberts and Maciejewski, 1992***; ***Mukherjee, 1995***). Meanwhile, others have focused on solving for the joint-space trajectories with optimization-based methods aimed at minimizing some cost function (***Biess et al., 2007***; ***Nguyen et al., 2021***; ***Flash et al., 2019***; ***Wolpert et al., 1995***), or using dimensionality reduction methods enabling a search for coupled or correlated DoFs (***Gritsenko et al., 2016***; ***Schröder et al., 2014***).

There also has been research using optimization and computational methods as a means to explain neural motion planning. For instance, the minimum-jerk model (***Flash and Hogan, 1985***) was developed to account for the kinematic characteristics of observed human hand trajectories, assuming that movement is coordinated to achieve optimal smoothness of the end-effector trajectory. Other studies have developed different models based on the minimization of alternative cost functions, such as the minimum torque-change (***Uno et al., 1989***) and the minimum variance (***Harris and Wolpert, 1998***) models.

More recently, other elaborate optimization models were developed aiming at integrating multiple costs at different levels of the motor hierarchy in order to provide a complete solution. For example, many models (***Denny Fu et al., 2013***; ***Menner et al., 2021***; ***Isableu et al., 2016***; ***Mombaur et al., 2010***; ***Berret et al., 2011***; ***Oguz et al., 2018***; ***Sharif Razavian et al., 2015***; ***Scott, 2004***) have emphasized the significance of optimal feedback selection for successful generation of multi-joint movements, based on the optimal feedback control framework developed by ***Todorov and Jordan (2002***), and some others have exploited machine learning techniques (***Srisuk et al., 2017***; ***Liu and Liu, 2020***).

On the other hand, a different set of approaches to the inverse kinematics problem have exploited dimensionality reduction techniques. The idea behind this class of approaches is that appropriate motor commands might occupy only a subspace or manifolds of lower dimensionality within the high-dimensional space of possible solutions, involving different combinations of motor variables and control inputs (***Flash and Bizzi, 2016***; ***d’Avella et al., 2015***). In that case, correlations between the DoFs can be identified, and these correlations quantitatively reflect coordination patterns responsible for effectively lowering the dimensionality of the motor representation.

The law of intersegmental coordination is a good example of such an innate dimensionality reduction solution. Motor control scientists have described a behavioral phenomenon whereby the absolute elevation angles of the leg during locomotion essentially covary on a plane, known as the intersegmental plane of coordination, and can be represented by two degrees of freedom, instead of the three existing joint angles (hip, thigh, and shank) (***Bianchi et al., 1998***; ***Borghese et al., 1996***). The absolute elevation angles describe the orientation of the leg segments with respect to gravity as opposed to the anatomical angles that describe the orientation of one leg segment with respect to an adjacent segment. The observed covariation plane can also provide context that can be useful for understanding healthy and pathological gait patterns (***Israeli-Korn et al., 2019***; ***Gueugnon et al., 2019***; ***Martino et al., 2014***). In particular, the observations described by this law of intersegmental covariation are only obtained if the joint angles are represented in terms of absolute coordinates in the sagittal plane, which is a finding that indicates the importance of identifying which reference frames subserve the representation of motor commands during human gait.

In recent years, numerous dimensionality reduction methodologies have been applied to kinematic and dynamics data obtained from studies of the motor system, resulting in different definitions of motor variables as well as units of action and motor primitives. These are usually selected based on geometrical considerations, statistical likelihoods, information theory concepts, etc. Specifically, in this work we have reformulated dimensionality reduction by using a blind source separation family of solutions. In particular, we applied a dimensionality reduction method called FADA (Fourier-based Anechoic Demixing Algorithm) developed by ***Omlor and Giese (2007***); ***Chiovetto et al. (2016***) based on anechoic demixing to derive the basis functions (also called sources) that underly movement patterns. Specifically, we consider whether it may be possible to use the same single set of basis functions to represent both joint and task position variables. This would then provide a hypothesis for how the nervous system may solve the inverse kinematics problem. Details of this method are presented in the Background in the Dimensionality Reduction section.

Additionally, in the Background we include a description of the arm kinematic model presented by ***Soechting (1982***), on which our study is based. Finally, in this work we focus on kinematic and mathematical analyses to determine which reference frames might allow for a unique set of basis functions to represent both joint and task spaces. Particularly, we examine the joint-space in an anatomical reference frame versus an absolute reference frame. See the Background for a longer discussion about the relevance and importance of different reference frames and representations.

## Background

The goal of this work is to help solve the inverse kinematics problem for arm movements using dimensionality reduction to identify basis functions that may be used by the CNS for movement coordination. An important element to consider in a given solution, is what coordinate system any solution is represented in, since this can provide insight into how the nervous system resolves redundancy. Additionally, it is important to carefully select the arm model used in obtaining results, as this may affect the solution and some arm models may allow for more impactful results. Finally, the specific method of dimensionality reduction used for identifying motor primitives or basis functions is important as some methods may produce more meaningful, generalizable results. These topics are discussed in this section.

### Reference Frames & Representations

A multi-joint system can be described by its configuration at each instant in time; however, this depends on the coordinate system used for its spatial representation. The various possible coordinate systems are termed *generalized coordinates*, which are sets of variables that uniquely and compactly define the configuration of the system (coordinates in the configuration space). Figure 1 exemplifies two possible coordinate systems for a simple kinematic chain. Theoretically one could define an infinite number of coordinate system *representations*, but it turns out that these representations can be grouped into specific classes. Two representations in the same class are equivalent if a rigid/linear transformation between them exists. Therefore, two questions should be asked for a given system: **a)** can the controlled representation be identified? **b)** what are the reasons for choosing one representation over another?

**Figure 1:**
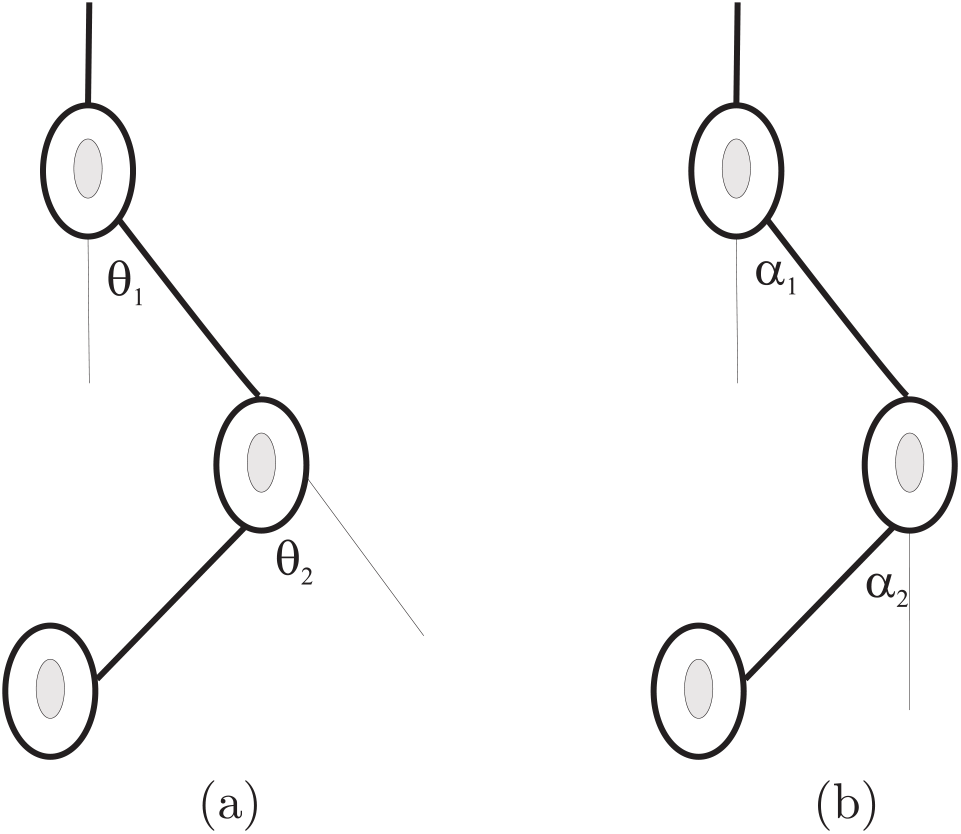
Generalized Coordinates: Two different representations that describe that exact same configuration: a) Anatomical or relative joint angles, b) Absolute or Elevation angles.

The question of representation is critical in motor control. Many studies have focused on which variables (kinematic, dynamic, etc.) are represented and controlled by the CNS for movement planning and execution. Over the years, this question has attracted considerable attention and among other studies, Soechting and Flanders and colleagues have extensively studied this issue. Specifically, in ***Soechting and Ross (1984***) they conducted several psychophysical studies aimed at examining alternative representations to determine which particular representation subserves the control of human arm posture and movement. Their experiments consisted of a *matching task* in which one arm was set to a given joint angle by the experimenter and the subject was asked to match this joint angle with their other arm. Movement in the matching arm was constrained to the degree of freedom being investigated. Their working hypothesis was that the “natural” coordinate representation of joint angles would have the lowest standard deviation in the difference between the joint angles of the two limbs.

In ***Soechting (1982***), the authors investigated three different coordinate systems for the shoulder orientation and two different systems for the elbow, and according to their psychophysical results discovered that the sense of limb orientation appears to be expressed best by the coordinate system illustrated in Figure 2b, picking (*θ, η, α, β*) to represent the arm configuration. It should be noted that rather than using relative orientation, such as defining the forearm joint angle with respect to the upper arm (*ϕ*), these angles describe the limb orientation relative to an absolute frame of reference.

**Figure 2:**
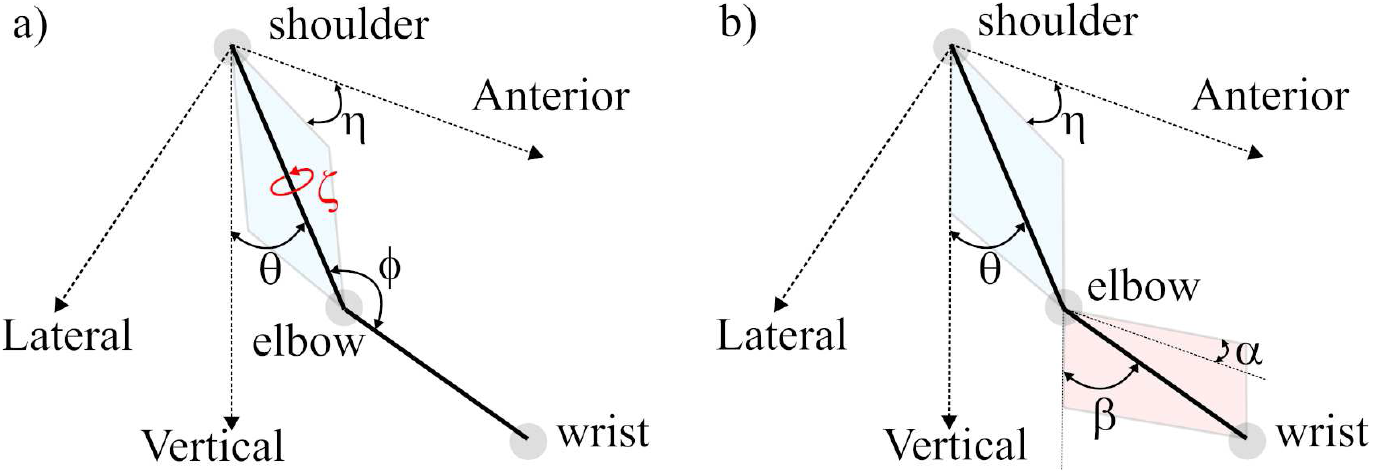
Arm model. In a) the first representation (*θ, η, ζ, ϕ*) combining absolute and internal reference frames is shown and in b) the second representation (*θ, η, β, α*) with a completely external reference frame is shown. The plane of the upper arm is shown in blue and the plane of the forearm is shown in pink.

***Borghese et al. (1996***) investigated coordination patterns of leg segments and joints during locomotion. Importantly they examined leg joint coordination in two sets: anatomical (relative) joint angles and elevation angles (absolute). In their paper they claim that these two representations form different classes and no simple transformation exists between them. The hip angle can be used to illustrate this point. The thigh elevation angle is the angle between the thigh and the vertical, and the hip angle is the angle between the thigh and the torso. Thus the hip angle cannot be reconstructed unless the torso is assumed to be vertical. Therefore, the transformation between coordinate frames without torso orientation is not trivial.

Furthermore, it was shown in multiple studies (***Borghese et al., 1996***; ***Bianchi et al., 1998***; ***Barliya et al., 2009***) that elevation angles are well behaved, have highly sinusoidal properties and when plotted with respect to each other, they lie on a plane during the gait cycle. On the other hand, the time course of the anatomical angles is much more variable, including intercycle variability. Additionally, ***Soechting and Ross (1984***) also found that subjects were best able to match joint angles of their right and left arms when they were measured relative to the vertical axes and the sagittal plane.

In particular, these experiments identified yaw and elevation angles as the selected spatial coordinate system for the perception of arm orientation. Studies have shown that target location is initially defined in a reference frame centered at the eyes and the origin of this reference frame is shifted toward the shoulder during the neural processing for targeted arm movements (***Soechting et al., 1990b***). Then, in this shoulder-centered frame of reference, target location is defined by three parameters: distance, elevation and azimuth (***Soechting and Flanders, 1989***). Moreover it was suggested by ***Flanders and Soechting (1990***) that there exists a linear transformation involving two separate *channels*: Arm elevation is computed from target distance and elevation, and arm yaw is computed from target azimuth.

In another study, ***Ivanenko et al. (2007***) suggested that there might be independent control of parameters in spherical representation of the end effector. In particular, they used PCA applied to elevation angle data and found a strong correlation between the PCA-derived basis functions of absolute angle basis functions and those of the polar and radial representatons.

These studies have all contributed to our understanding of how the nervous system represents and coordinates movements. For instance, it seems that the nervous system has some preference for an absolute frame of reference, and yaw and elevation angles seem to be keys to joint coordination. The validation of these insights is essential. To help validate these coordinate frame insights, in this work FADA will be used to find sources (“basis functions”) in both joint-space, represented by both the relative/anatomical and the absolute/orientation angles, and in task-space, represented by the end-point variables. Additionally, there still remain open questions related to the coordination of movement. Importantly, the intersegmental coordination of the arm has been neglected for the most part in the literature, which is a primary focus of this study.

### Arm Model

To model the arm we first can consider that an unconstrained rigid object in 3D space has six DoFs in total: three translational and three rotational. However, when there are multiple linked segments the number of DoFs is reduced due to kinematic constraints. For example, assuming no translational movements at the shoulder (glenohumeral joint), the upper arm has three DoFs, which can be modeled by a ball-and-socket joint allowing flexion/extension, abduction/adduction and internal/external rotations. The forearm, modeled as a hinge joint, adds two more DoFs: flexion/extension, and pronation/supination. Thus, there are a total of five DoFs for the simplified human upper limb (more if we consider the wrist and fingers). However, in the current study, we used a simpler model for the arm (i.e., not including elbow pronation/supination), with two rigid links joined at the elbow joint, ending with only 4-DoFs.

Specifically, our arm model and the angles we used as defined in ***Soechting et al. (1995***), is illustrated in Figure 2. In this model, the first angular rotation (*η*) is about the *Z*-axis and determines the yaw angle. The second angular rotation (*θ*) is about an axis perpendicular to the arm plane (this is the plane spanned by the vectors of the upper-arm and forearm, the arm plane is the lateral *X*-axis if there is zero yaw) and determines the arm’s elevation. The third angular rotation (*ζ*) is about the humeral axis. This rotation does not change the elbow location but does affect the wrist location in space and the arm plane. Finally, *ϕ* is the angle of flexion of the forearm, *ϕ* = 180^°^ corresponding to full extension.

It should be noted that Figure 2b contains two more angles: *β*, the angular elevation of the forearm, and *α*, the forearm yaw angle (just like *η* for the upper-arm). Thus, two sets of variables, or representations, can be devised to fully describe the arm configuration. The first representation (*θ, η, ζ, ϕ*) combines external (absolute) and internal (relative) reference frames (see Fig. 2a). In this case *θ* represents the upper-arm elevation angle, *η* is the azimuth angle, *ζ*the humeral rotation angle, and the elbow flexion-extension angle is given by *ϕ*. The second representation (*θ, η, β, α*) is given in a completely external frame of reference (see Fig. 2b). Here, *β* and *α*, denote the elevation and azimuth of the upper-arm, respectively. We term the first representation *anatomical* or *relative*, and the second is termed *absolute* or *external*.

### Dimensionality Reduction

Movement data (such as joint angles or muscle activations) generally has high dimensionality. However, regardless of the level of complexity, every arm movement is ultimately mapped to three Euclidean coordinates describing the hand position. Therefore, a set of correlations or a coordination pattern must exist in these higher motor levels that constrains any seemingly excess DoFs. The challenge is to identify these correlations.

One of the most well known methods for identifying coordination patterns is the Principal Component Analysis (PCA) method. This method involves a mathematical procedure that maps possibly correlated variables onto a small set of *uncorrelated* variables called principal components. The basic underlying assumption for PCA is that the observed data (*x*_*i*_(*t*)) can be modeled as a linear combination of orthonormal basis functions (*s*_*j*_), the vectors of which are the eigenvector of the covariance matrix representing the data, with time dependent mixing coefficients for the PCs.

Another well known method for dimensionality reduction is independent component analysis (ICA). In contrast to PCA the underlying model assumed by ICA uses a time dependent basis functions *s*_*j*_(*t*) with constant mixing coefficients. More importantly, here the basis is required to be independent, rather than uncorrelated as for PCA, which is a stricter constraint. ICA has classically been used as a solution to the *cocktail-party problem* (***Cherry, 1953***). This problem is a special case of the *blind source separation* (BSS) problem.

An alternate version of the BSS problem is anechoic blind source separation. Similarly to ICA, this method aims to solve the BSS problem, while allowing for the addition of time delays between the sources

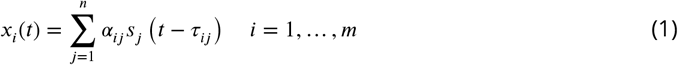

The model expressed in (1) appears in acoustic equations where a reverberation-free environment is modeled, i.e. the sensors only receive attenuated sounds with different arrival times. Thus mixtures of the form (1) have been termed *anechoic mixtures*. A solution for a system assuming a non-linear model such as described in (1) was proposed by ***Omlor and Giese (2011***). The solution assumes that the sources are uncorrelated and uses properties of the stochastic Wigner-Ville spectrum (WVS) (***Matz and Hlawatsch, 2003***). The solution was obtained by representing the signals in the time-frequency domain using the Wigner-Ville transform, which is defined by

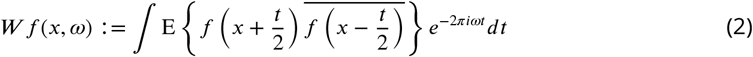

where E denotes the expected value and the bar denotes the complex conjugate. Applying this transformation to Eq. (1) and exploiting the (approximate) independence of the sources, one obtains

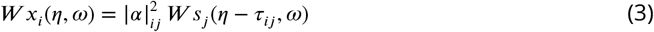

Assuming that the observed data coincide with the mean of the distribution (*x*_*j*_ ≈ *E*(*x*_*j*_)) one can compute the zero moment of Eq. (3) and obtain the following two equations:

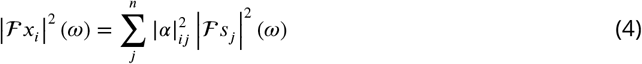

and

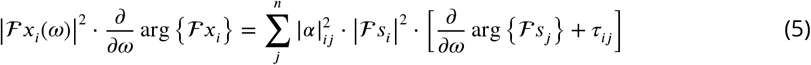

where ℱ is the Fourier transform operator. Non-negative ICA was used to solve Eq. 4, the results of which are used in Eq. 5 in order to extract the corresponding time delays. The latter was solved in an iterative manner as detailed in the above mentioned papers. Therefore, the solution of the above system of equations results in the requested set of sources (*s*_*j*_) and the corresponding time delays (*τ*_*ij*_).

This method models high-dimensional signals as a linear superposition of a small set of source functions, which can have additional fixed temporal delays. The solution results in a set of sources for each space to which it was applied. Then, for each movement type the proper mixing weights and time delays were determined. Thus, this approach can allow for a simple computational solution to the redundancy problem, based on the assumption that the same basis functions underly the joint and task spaces.

Here, a new approach to resolving the redundancy problem was suggested that exploits dimensionality reduction called FADA (Fourier-based Anechoic Demixing Algorithm) (***Chiovetto et al., 2016***), which is based on a computationally efficient form of anechoic demixing algorithm for the analysis of band limited signals ***Omlor and Giese (2007***). This was originally derived from a more general anechoic demixing algorithm developed by ***Omlor and Giese (2011***). FADA is a highly efficient dimensionality reduction approach and an alternative way to characterize motor primitives based on the idea that they express invariance across time. In particular, in this work we use FADA to produce basis functions (or sources) for either the task or joint spaces. This can help researchers gain a greater understanding of how the CNS possibly generates desired task space trajectories with appropriate movements of redundant joint DoFs. Specifically, FADA provides basic temporal patterns or functions *s*_*p*_(*t*) which are combined or superposed to reconstruct a set of temporal signals. Hence, the temporal decomposition is mathematically described as:

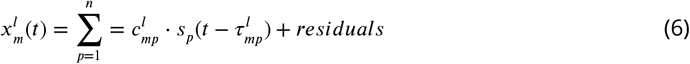

In this equation 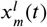 is the value of the m-th DoF at time t in trial number l, and the corresponding scalar mixing weights 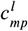 change between trials of different types (experimental conditions), P signifies the total number of temporal primitives. This model also allows for time shifts between the temporal basis functions for different DoFs, which are captured by the variables 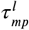. The time delays and linear mixing weights are typically assumed to vary over trials, while it is assumed that the basis functions *s*_*p*_(*t*) are invariant ***Chiovetto et al. (2016***). Details of the iterative FADA algorithm are described in more detail in the Source Separation section of the methods.

## Materials and Methods

### Motion Capture & Data Processing

Arm movements were recorded for fifteen subjects (aged 28-35) who volunteered for the experiment. None reported previous hand injuries and all gave their informed consent prior to their inclusion in the study.

Subjects were grouped based on the three motion sets they completed: one group had four subjects who each conducted all motion types (ALL), one group had six subjects who only completed the figure eight motion (FE), and one group had 5 subjects who only completed the planar ellipse motion (PE), though at different orientations in space. For future reference these groups will be referred to as ALL, FE, and PE, respectively.

Subjects were instructed to freely draw these series of shapes in three-dimensional (3D) space repetitively at a comfortable pace with their dominant arm. Ten paths were drawn by the subjects in the ALL group: planar ellipse (PE), vertical ellipse drawn on the sagittal plane (PE-V), ellipse drawn on the frontal plane (PE-F), planar ellipse drawn on a plane rotated 45^°^ off the sagittal plane (PE-45V), ellipse drawn on a plane rotated 45^°^ off the frontal plane (PE-45F), horizontal figure-eight (FE), vertical figure-eight drawn on the sagittal plane (FE-V), figure-eight drawn on the frontal plane (FE-F), bent-ellipse (BE), double bent-ellipse (DBE) and up-down movements (UD). The models of the shapes are presented in Figure 3. The drawing instructions did not involve the experimenter demonstrating the movements to avoid biasing the subjects towards a specific behavior. Therefore, we prepared wire frames models for the different shapes and showed them to the subjects before the recordings of each shape began.

**Figure 3:**
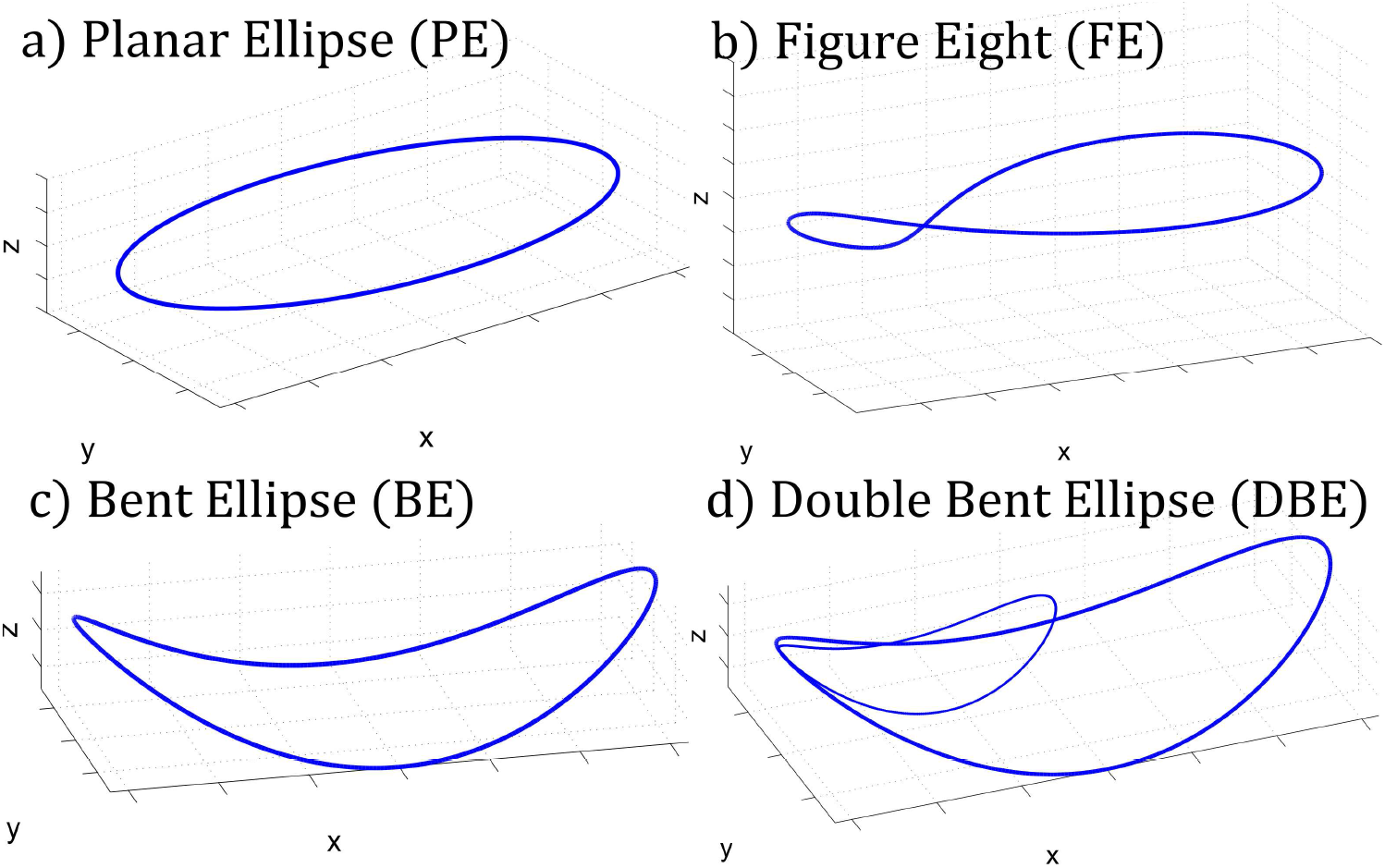
Wire frame models of the recorded shapes, as shown to subjects prior to data collection. These were the main shapes, but the FE and PE shapes were also collected at different orientations.

Subjects were seated on a wooden chair with a high rigid back rest. Movements of the ALL group were recorded using a Polhemus “Liberty” electromagnetic spatial tracking system where sensor positions and orientations were collected at 240 Hz and preliminary experiments were conducted to measure the accuracy of the tracking system, and the error was found to be at most *θ*.3mm. Eight sensors were positioned on the arm: one on the wrist, two on the forearm so as not to be co-linear with the wrist marker, two on the upper-arm, one on the shoulder (again, ensuring these three sensors are not co-linear), one on the chest near the collarbone, and one on the chair acting as reference. The tracing of each shape was recorded continuously for 20 seconds. For the ALL group two 20-second trials were carried out for each shape per subject. Thus, the data base of movement for the ALL group consists of 4 (subjects) × 10 (shapes) × 2 (trials per subject) = 80 recordings (see example PE data in Fig. 4).

**Figure 4:**
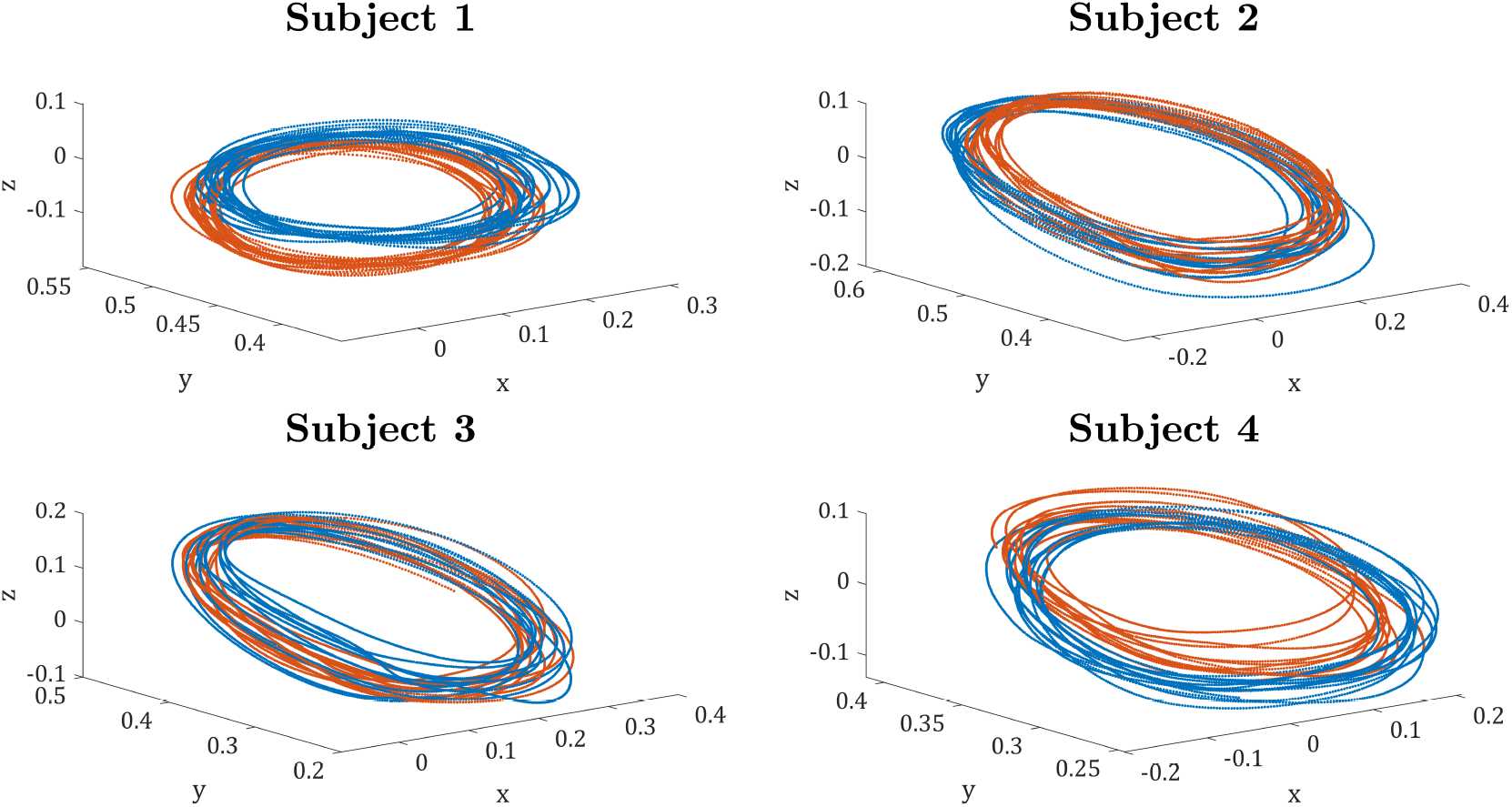
Data of the planar ellipse (PE) shape for the four subjects from the ALL group. Each subject completed two 20-second trials, shown in 1) blue, 2) orange. Note variability in the traces for each subject.

Meanwhile for the PE and FE groups, movements were recorded using a Vicon IR motion capture system. We analyzed the data using a simplified arm model in which the end-effector is at the center of the wrist, the task space has 3 DOFs and the configuration space has 4 DOFs (3 in the shoulder and one in the elbow). In each trial subjects had to draw the shape for 25 seconds with their healthy, dominant right hands.

The data were interpolated for missing samples, then approximated by smooth analytic curves (to allow high-order differentiation). The continuous data were then segmented into individual repetitions. Though the use of two different motion tracking systems could produce slight differences between the subject groups, we believe it shouldn’t significantly impact the results of our source reference frame or timing comparisons.

### Joint Angle Extraction

Joint angle calculation is not a trivial task. First, one has to identify the centers of rotation (COR) of the arm located at the shoulder and elbow, and in some cases it is not possible or simple to place a marker exactly on the joint center. This problem is crucial to the study of human movement and biomechanics, including the calculation of joint angles. Many approaches have been used in the literature to try to address this problem. We have chosen to use an extension of the technique that was described in ***Gamage and Lasenby (2002***).

### Source Separation

While there are many methods of performing source separation such as PCA and ICA, in this work we used the FADA algorithm as first presented in ***Chiovetto et al. (2016***). This algorithm can allow for source separation without constraints but is also flexible to allow for parameter non-negativity or other constraints.

Specifically, this iterative algorithm is as follows:

1. Non-negative ICA is used to solve the equation

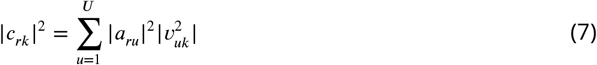

with r = 0, 1, … R and k = 0, 1, … K. Non-negative matrix factorization [34, 41] can also be used to solve this equation instead of ICA. Specifically here, the mixing weights are represented by *α*_*ru*_, while *c*_*rk*_ and *v*_*uk*_ represent the coefficients of the Fourier expansion of the *r*-th joint angle and *u*-th temporal source signals, respectively.
2. To obtain the phase shifts, the Fourier cofficients are then updated by solving the nonlinear least square equation

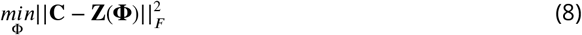

where *F* is the Frobenius norm and **C** and **z** are given as **C**_*rk*_ = *c*_*rk*_ and

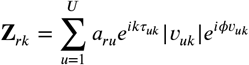
3. The weights *α*_*ru*_ and delays *τ*_*ru*_ are then identified for each signal *y*_*r*_(*t*), by minimizing the cost function:

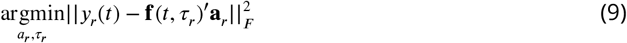

Here the source functions *f*_*u*_(*t*) are kept constant (and assumed to be uncorrelated), and it is assumed the time delays are independent. Specifically, the vector **f** (*t, τ*_*r*_) is the concatenation of these functions including the time shift associated with each DoF *τ*.

### Inverse Kinematics

In this section we show that given two arbitrary spaces (possibly of different dimensionalities) that share a similar set of basis functions, one can transform between the spaces using the basis functions as mediators. This idea was originally proposed based on some of our previously obtained results. Specifically, we would like to show that there is a mechanism to determine the joint-space trajectory given a desired trajectory in task-space, and assuming that the task-space and joint-space movements share the same set of sources. We begin by formally defining the problem itself.

#### Problem Statement

As mentioned, we assume a nonlinear model underlying both the joint and task-spaces. Thus, the basis functions can also be shifted in time. Let the *shift operator U*_*λ*_, applied on a periodic function *f*, be defined as follows

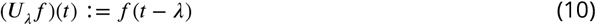

The shift operator *U* essentially delays the function *f*(*t*) by *λ* in a circular periodic manner. This is actually achieved through the *convolution* operation

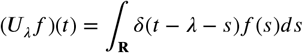

where *δ*(*x*) is the Dirac delta function. The above definition is for the one dimensional case, and next we define a more complex operator for the multidimensional case.

Let 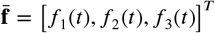 be a vector of periodic functions (the extracted sources). We define the *shift operator matrix* as follows

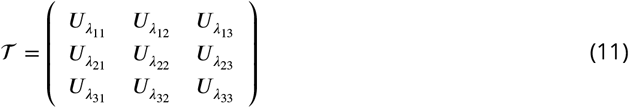

where 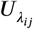 are the standard shift operators as defined above (Eq.10). Then, applying ℱ on 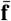 we get,

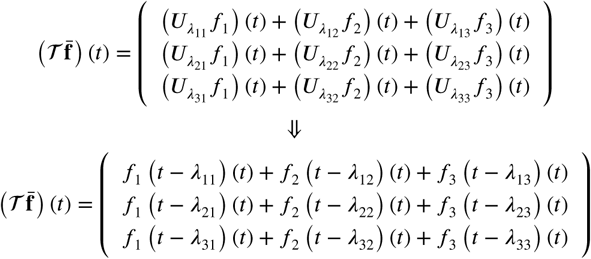

Thus, the *problem* is to define the inverse of 𝒯 (𝒯 ^−1^) such that if

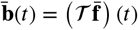

then applying 𝒯 ^−1^ will result in

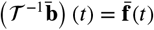

The problem can be slightly extended to the *weighted* version of the operator 𝒯. Let,

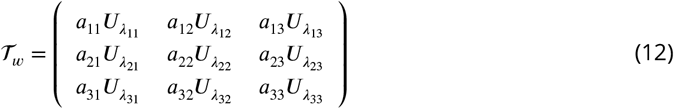

where *a*_*ij*_ are weights. Applying 𝒯_*w*_ on 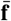 is just

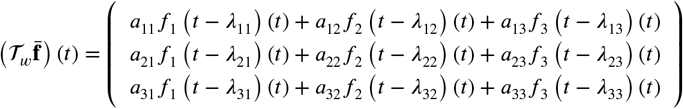

So similarly to the previo us inverse, 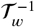 should be defined. In effect, having such an inverse applied on, say the task-space trajectory, we will be able to obtain the sources in return.

#### Simplified Case (without time delays)

We begin with a simpler version of the above problem, in which the sources are not shifted in time, i.e., *λ*_*ij*_ = *θ*. Let *s* = (*s*_1_, *s*_2_, *s*_3_) be the vector of the extracted sources. The sources are assumed to be identical for both spaces, thus the joint angles, *q*, can be expressed as

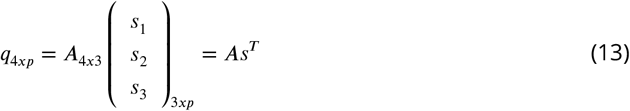

Similarly, using the same sources the Euclidean coordinates, *x*, can be expressed as

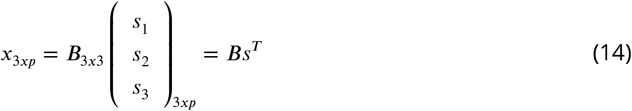

where A and B are the matrices of amplitudes associated with the sources, and p is the number of points in the path.

In order to attend to the first case of a direct mapping between joint and task-spaces through the forward kinematics (*f*_*i*_ : **Q** → **X**), observe that the dimension of B is 3 × 3. Thus, in a well-behaved case and relying on the independence of the sources, B can be regarded to be invertible and therefore

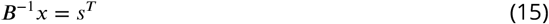

Substituting s from Eq.(15) into Eq.(13) we obtain:

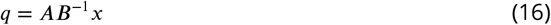

This is an inverse mapping of positional variables (not velocities). Note that although the sources s were the original mediators, they do not appear in the transformation and only their amplitudes are required.

The second case, involving the mapping of instantaneous positions is a bit more complicated. Specifically, the mapping between joint and end-effector velocities (or instantaneous position) for forward kinematics is given by

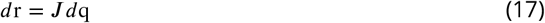

with *J* as the Jacobian of the forward kinematics. Using Eq.(17), Eq.(13) can be re-written as follows:

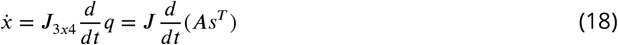

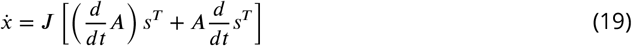

since

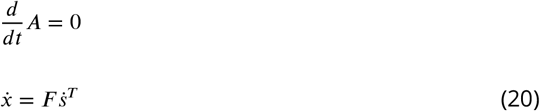

where *F* = *J A* is of dimensions 3 × 3 and potentially invertible.

This takes the form of a coordinate transformation into a space that can be uniquely inverted to the task-space. Thus the first derivative of the joint angles, 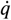, can be expressed as a linear combination of a new set of sources 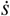. We find this has the form

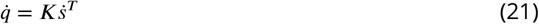

where *K* is a matrix containing the coefficients of 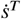, and *K* = *AB*^−1^*F*.

So far though, the solution is only for the degenerate case where time delays are ignored. This is not a realistic state, and it will be remedied in the next section.

#### Inverse Solution

Next, we address how to integrate the required time shifts into the inverse transform, using a circular shift operator.

A convolution operator *A* is formally defined as

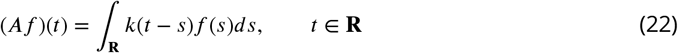

The function *k* in this context is referred to as the *convolution kernel* of *A*. Let ℱ : *L*^2^(**R**) → *L*^2^(**R**) denote the *Fourier transform*,

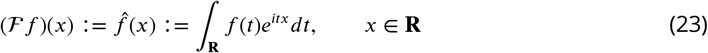

and let ℱ^−1^ : *L*^2^(**R**) → *L*^2^(**R**) be the inverse of ℱ, given by

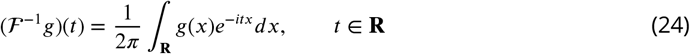

The operator (22) can formally be written in the form

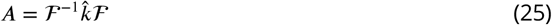

or equivalently, 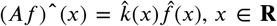. The function 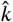 is called the *symbol* of the operator (22), (25).

Now we consider a special class of convolution operators on *L*^2^(**R**).

Fix *λ* ∈ **R** and let *U*_*λ*_, a bounded linear operator^1^, be the *shift operator* defined by

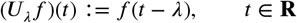

Since

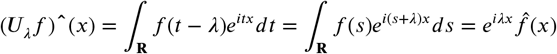

Using the Dirac delta function *δ*(*t*), we have

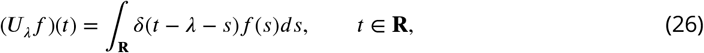

which shows that *U*_*λ*_ is a convolution by the kernel *δ*(*t* − *λ*). The convolution operator is often expressed as a binary operation between a function *f* and a kernel *k* denoted by

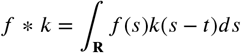

Therefore, according to the above definitions and properties, it is clear that the convolution commutes with the shift operator (translation), that is

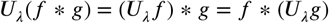

To connect this to our problem, we assume that each signal can be represented a follows:

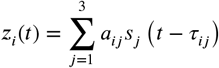

Let us assume that a similar set of sources underlie the joint-space and task-space and representing the joints and hand behavior by (13) and (14), respectively. Consider first the representation of a one dimensional signal in terms of the convolution and shift operators,

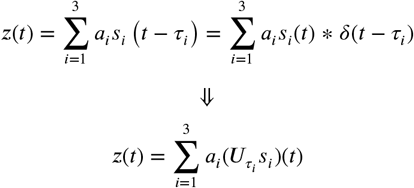

where 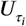 is the shift operator described in (26). We can now extend the one dimensional case to a multidimensional space.

Let

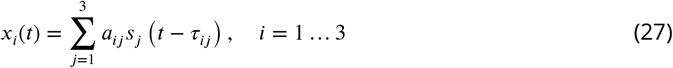

where *x*_*i*_ are the Euclidean coordinates of the hand. Thus, the hand coordinates can be represented by a mixture (sum) of the weighted (*a*_*ij*_) and delayed (by *τ*_*ij*_) sources (*s*_*j*_(*t*)). Equation (27) can be expressed differently using the convolution operator, as follows

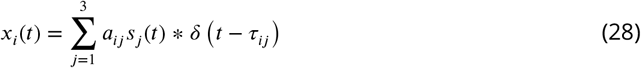

where *δ*(*t*) is the Dirac delta function, and the symbol ∗ stands for convolution. Now, applying the Fourier transform on both sides of equation (28), we obtain

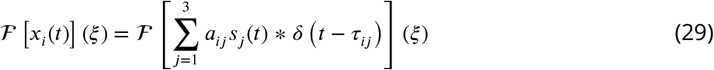

where ℱ is the fourier transform defined by Eq.(23). Due to the linearity of the Fourier transform, equation (29) can be rewritten as

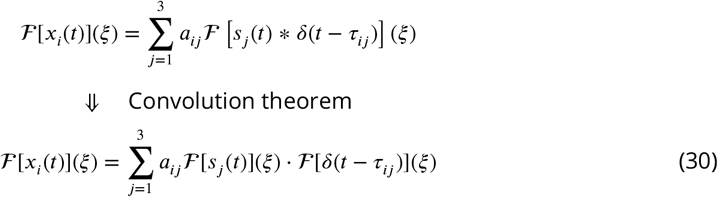

The Fourier transform of the delta function is

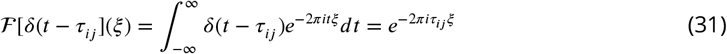

Substituting equation (31) into (30) we get,

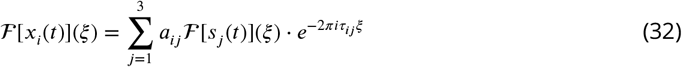

The system of equations (32) is the frequency domain version of the temporal domain system (27). In this domain, the inverse is more easily expressed.

Let the matrix

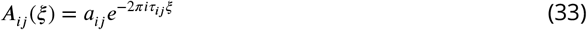

be the matrix that contains the information of the mixing coefficients of the sources and the their corresponding time-delays. Then, system (32) can be expressed in matrix form as

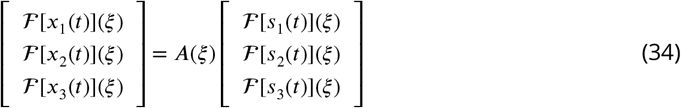

where *A*(*ξ*) = [*A*_*ij*_(*ξ*)] is a 3 × 3 matrix. Then, by inverting *A* we get,

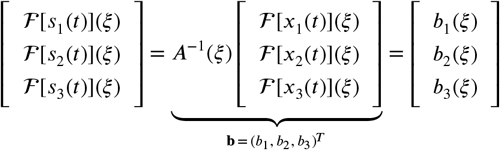

Applying the inverse Fourier transform on both sides of the above system leads to

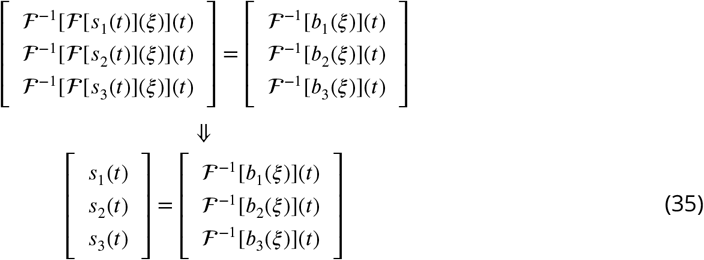

and we have the solution for the inverse problem. The solution here, is having the sources themselves extracted as a function of the task-space.

An important note to mention is that the solution above assumes that the matrix *A*(*ξ*) is invertible. The matrix *A* might not be always invertible, but its properties can be analyzed and this has been considered and presented in Appendix **A**. This analysis ultimately showed that *A* is invertible if and only if |*A*| ≠ 0.

Thus far, we have shown that provided two spaces share a basis it is possible to go back and forth between the spaces. In fact, the proof itself illustrates this methodology. Furthermore, the CNS could hypothetically utilize this to solve the inverse kinematics problem using the basis functions as mediators between the spaces. However, there are some knowledge gaps left to fill. Assume that we are planning a movement in task-space, and would like to produce the corresponding joint-space trajectory. The results of the previous section state that the joint-space trajectory can be expressed by:

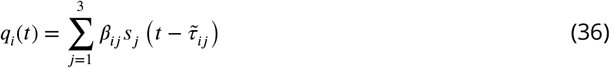

where *β*_*ij*_ and 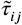 are the appropriate weights and time shifts, respectively. Furthermore, similarly to the process in the previous section which concluded in Eq.(33), the time delays 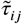 can be pulled out of the sources and be represented. Then, *sj* can be replaced with Eq.(35), thus *q*_*i*_(*t*) is expressed in terms of *x*_*i*_ (*t*) (the task-space)

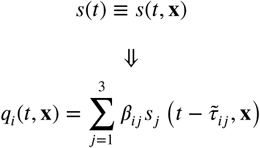

Indeed, the above solution shows how the joint-space 𝒬(*t*) trajectories can be expressed in terms of the task-space trajectories 𝒳(*t*). However, the parameters *β*_*ij*_ and 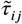 are not an outcome of the above process and need to be determined. In the next section, we will investigate the sources and their phase shifts, which will provide us with clues on how the time delays are determined.

### Source Analysis

Each shape is described by a set of task space sources and a set of joint space sources. These sources can be compared in two major ways: a) by analysing how similar the source shapes are to one another and b) by analyzing how the sources are shifted in time, or phase, between each other.

#### Source Shape Analysis

The joint-space sources (both representations) were compared with those of the task-space both visually and numerically (see Figure 8 for description). To aid visual comparison between the sources they were normalized using their ℒ_2_ norms. Then, the cross-correlation was computed between the different sources (each source in joint-space was cross-correlated with all other sources in the task-space) to both find a match between sources and shift them in time with respect to each other to overlay the sources in one plot. The cross-correlation between two curves *f* and *g* is defined as

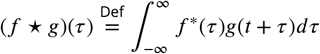

where *f*^∗^ denotes the complex conjugate of *f*.

The cross-correlation similarity measure index (similarity index or SI for short), defined as the maximum correlation value between two signals, was compared between each task-space source and its joint-space counterpart, given by the anatomical and absolute orientation-angle sources. For very similar signals, values that are close to 1 will be achieved at zero (or close to zero) time lag.

We performed a two-way ANOVA test to determine if the differences between the task and joint space sources were statistically significant, while accounting for the differences due to subjects and selected movement shapes. The factors that were considered for our ANOVA were the angle type (absolute vs anatomical angle), the shape (i.e. FE vs PE etc) and the subject. Subject was considered a random factor and was nested within the shape group, as some subjects only completed movements of a single shape. The null hypothesis in this case was that there was no difference in the source correlations (i.e. neither set of joint space sources was more closely correlated to the task space sources). The results of this analysis would simply signify that there is a significant difference in the source correlations, but the magnitude and direction of that difference would need to be analyzed more carefully using other methods and post-hoc comparisons.

We also used the sign test to evaluate if there are statistically significant differences between sources. This is analogous to a non-parametric t-test, with no assumptions of a normal distribution. In particular, the sign test looks at whether one signal (or in this case, one source) is statistically larger or smaller than another. This is achieved by forming a vector where each entry indicates if the corresponding entry for one signal, i.e. source one, is smaller than the matching entry for signal two, in this case source two. The null hypothesis here is that the distributions are equal, and therefore there is an equal likehood for entries from one signal to be larger than entries in the other signal. The proportion of entries in the comparison vector that indicate one source being smaller than the other can be compared using using binomial distribution tables. For two matching sources the proportion would be 0.5.

So we use a normal approximation of the binomial distribution to calculate the z score. The z-score of the data could then be compared to the z-score of the null probability (we made sure that the estimated probability p and the size of the data n were consistent with n⋅p > 5 and n⋅(1-p) > 5.) to determine the p-value. Values of *p* < 0.05 were considered significant and indicated that the sources were not statistically equivalent.

#### Source Phase-Shift Analysis

Each movement cycle for an individual shape is distinct. This distinction becomes apparent when considering the weights and the time-delays of the sources that generate the movement. We per-formed an analysis to represent patterns of the time-delays (*τ*_*ij*_), assuming that these are not arbitrarily determined, and to expose correlations between the time-delays for different sources, i.e. phase-shifts. In particular, we will carry out the time-delay analysis for the absolute joint-space and task-space representations. Phase-shifts for 3D shapes and planar shapes will be separately considered.

#### Consider the following system

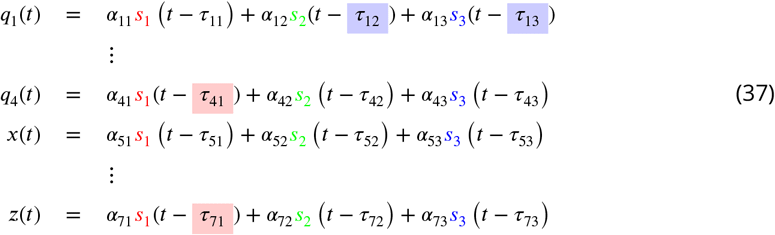

The phase-shifts can be analyzed across two levels. Firstly, phase-shifts between sources of the same degree of freedom were examined, i.e., *τ*_*ij*_ − *τ*_*ik*_ (see for example the blue blocks in Eq.(37): *τ*_12_ − *τ*_13_). Secondly, phase-shifts between different degrees of freedom in a single source were examined, i.e., *τ*_*ij*_ − *τ*_*kj*_ (for example, red blocks in Eq.(37): *τ*_41_ − *τ*_71_). Constant phase-shifts of the former type, if appearing, may imply coordination between different sources. One can think of the sources as movement generators, and thus constant phase-shifts would be the result of coordination at the level of these generators.

The second type of phase-shifts can appear due to coordination between different degrees of freedom, such as joint level coordination. The phase-shifts are computed as the absolute value difference between the sources’ phases, corrected to their relative percentage of one drawing cycle (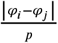, where *p* is the number of sample points in a movement cycle).

## Results

In this section we will show the results of the source separation for our data.

### Source Analysis Results

Sources were extracted from recorded data from all subjects. Visual inspection of these sources can provide insight into the significance of the anatomical and absolute angles.

#### Individual Source Examples

In Figure 5 example source results for each shape are shown for a typical subject in the ALL test group (for group definitions see methods section 3.1 on the motion capture). Meanwhile, Figures 6 and 7 present example source results for individual subjects from the PE and FE groups, respectively. Across all subject groups the absolute angle sources appear more similar to the end effector sources than the anatomical angle sources.

**Figure 5:**
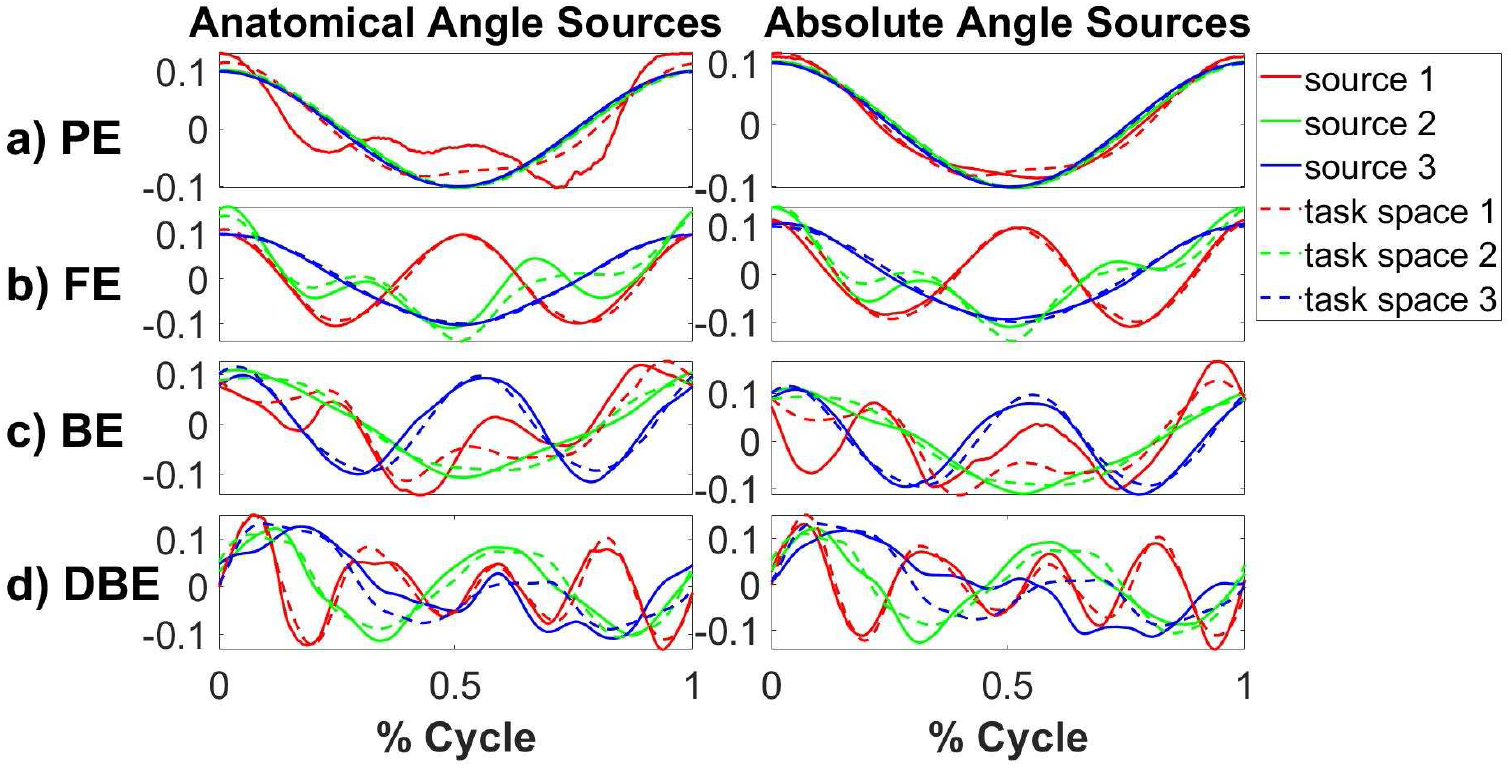
Sample source results from a typical subject from the ALL test group. Example sources are shown for the four shape types: a) PE or Planar Ellipse, b) FE or Figure Eight, c) BE or Bent Ellipse, d) DBE or Double Bent Ellipse. For each shape, sources based on the anatomical angles (shown on the left) and sources based on absolute angles (shown on the right) are compared to the end effector task space sources (see dashed lines). Using visual inspection alone it is clear that the absolute angle sources generally provide a better fit for the task space sources in this example, particularly for the PE and FE cases.

**Figure 6:**
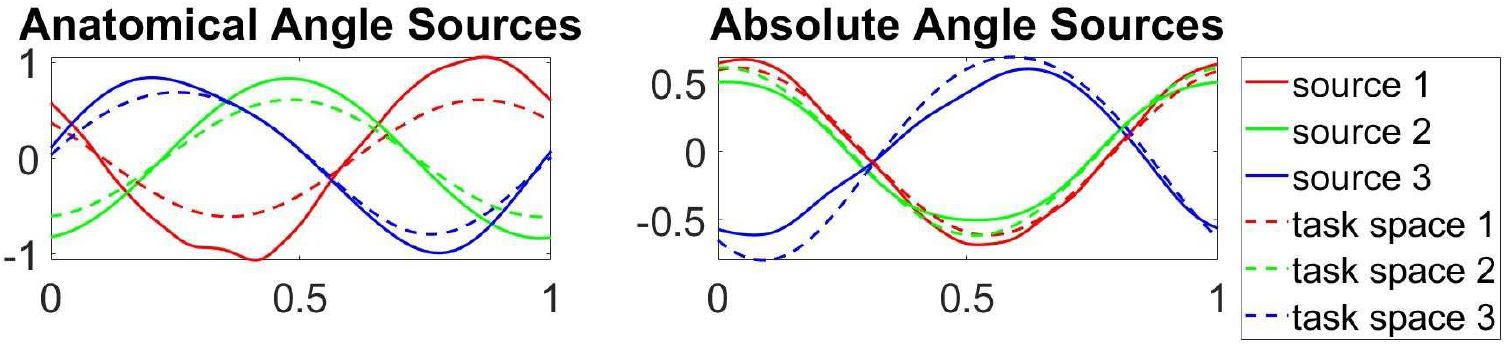
Sample source results for a typical subject from the PE test group. As shown in Figure x the anatomical angle sources are shown on the left and the absolute angle sources are shown on the right, both compared to the end effector task space sources (shown in dashed lines). Note that the shape is similar to that of the PE sources for the ALL subject. However, the sources are phase shifted compared to the consistent cosine graph shape of the ALL subject.

**Figure 7:**
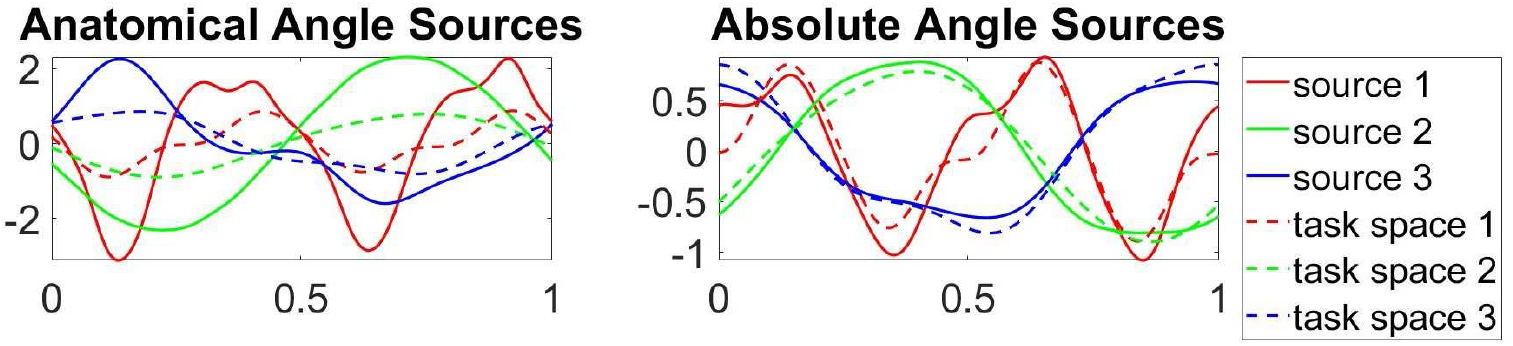
Sample source results for an FE test group subject. Note differences between the shapes of these sources and those for the FE shape of the ALL example subject.

Note that the source shapes are somewhat affected by the orientation of the arm movements during the experiment. For instance, the PE group were not constrained to draw ellipses on planes of precise orientation (and thus there was variability in the orientation of movements combined to find the sources), whereas the sources for the PE shapes from the ALL group were collected at precise orientations for which separate sources were computed. We believe this is the reason that there is a phase shift between sources of the PE group and the sources for the PE shape from the ALL group, even though the shape of the sources are similar (see Figure 7 and 6).

#### Overall Source Shape Results

Figures 8 through 10 show the sources from each recorded shape, as extracted from the recorded data for all subjects. For each shape, three sources were extracted for both joint representations as well as for the task-space and overlays of the task-space extracted sources are shown on top of the sources extracted from the joint-space (with the left column showing the anatomical sources and the right showing the absolute).

**Figure 8:**
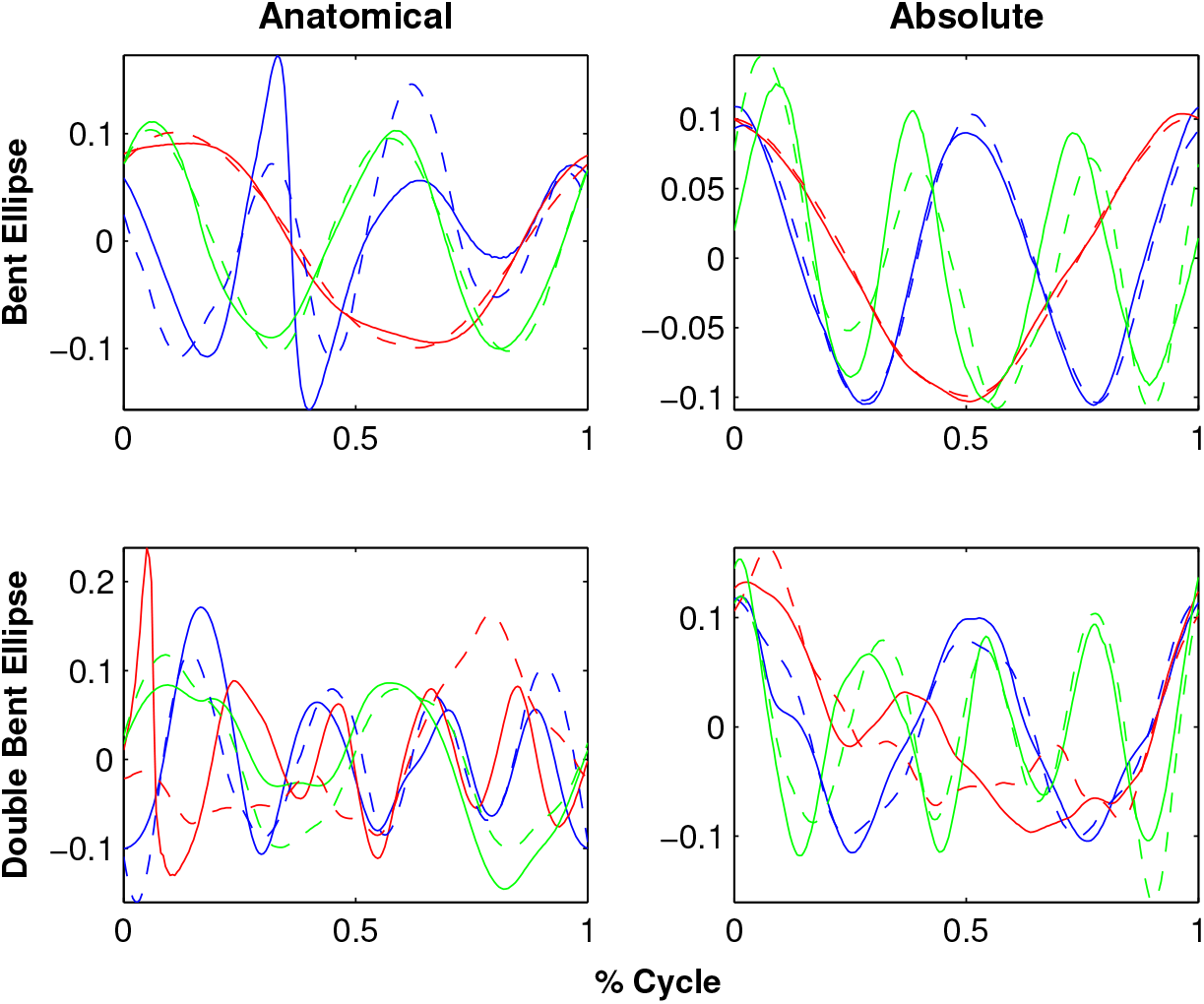
Extracted sources for the Bent Ellipse (top) and Double Bent Ellipse (bottom) shapes - The left column shows results for the joint-space using an anatomical representation, the right column shows results for the joint-space using an absolute representation. The task-space sources are overlaid over the sources of the anatomical angles representation (left column) and over the absolute angles representation (right column). Solid lines represent joint-space sources and dashed lines designate task-space sources.

Inspecting the overlays of the sources reveals a striking similarity between the task-space sources and the joint-space sources when represented by the absolute angles (orientation). There is a high degree of matching between these two sets of sources, and potentially one set of sources could be used to explain both the task-space and the joint-space when represented in its absolute orientation form.

This observation is supported by the cross-correlation similarity index values between the sources as they appear in Table 1. See the Source Shape Analysis methods section for explanation of how the similarity index is computed. This table shows the difference between the similarity index for the absolute angle sources compared to its counterpart end-effector (task-space) sources and the similarity index for the anatomical angle and end-effector sources. In particular, a distinction was made between the cases when the absolute angle sources had higher similarity to the end-effector sources than the anatomical angle sources (as signified by the positive values, shown in dark green). The results show that in most cases the absolute angle sources were more highly correlated with the task-space sources than were the anatomical angle sources, as signified by the values greater than zero (with higher values signifying greater differences in similarity indices between the two source types).

**Table 1:** Correlation Differences. The cross correlation was computed between the anatomical angle sources and the task space sources, and a second cross correlation was computed between the absolute angle sources and task space sources. The difference between these two cross correlation measures are presented here. The last column reports the differences between the means of the correlation indices. Positive values (shown in dark green) represent cases where the absolute angles have higher correlation with the task sources. Negative values (shown in light green) represent instances where the anatomical angles have higher correlation with the task sources.

| | $S_1$ | $S_2$ | $S_3$ | Mean |
| --- | --- | --- | --- | --- |
| FE X0 Y0 | 0.1010 | 0.026 | -0.16 | -0.011 |
| FE X90 Y0 | 0 | 0.2240 | 0.0070 | 0.770 |
| FE X0 Y90 | 0.007 | 0.011 | 0.085 | 0.034 |
| PE X0 Y0 | .001 | 0.054 | 0 | 0.018 |
| PE X45 Y0 | 0 | 0.009 | 0 | 0.004 |
| PE X90 Y0 | 0 | 0.001 | -.005 | -.002 |
| PE X0 Y45 | 0.615 | 0 | 0.034 | .217 |
| PE X0 Y90 | .36 | 0 | .008 | .122 |
| BE | .005 | .1460 | .006 | .052 |
| DBE | .099 | .71 | .114 | .308 |

As discussed in the methods section, the sign test and ANOVA statistical analyses were also conducted to ascertain statistically whether the anatomical or absolute sources were more similar to the end-effector coordinate sources.

The sign test used a normal approximation of the binomial distribution to calculate the z-score that would result for a probability of 0.5. This is equivalent to there being no difference between the two distributions. The results of the sign test yielded a p-value of p = 0.0018, which was statistically significant, assuming *α* = 0.05. Therefore, we conclude that there is a statistically significant difference between pairs of observations, which in this case represent the anatomical and absolute angle sources.

Meanwhile, the results of our two-way ANOVA are shown in Table 2. The p-values for “Type” (i.e. absolute vs anatomical angles) and for “Shape” (i.e. PE, FE, DBE, or BE) were 0.0054 and 2.95 ∗ 10^−^7 respectively, therefore these two factors are significant. Meanwhile, the random factor “Subject”, which is nested within the Shape factor, had a p-value of 0.2674, meaning this factor was not significant.

**Table 2:** Results of ANOVA. The p-values for the fixed reference frame and shape factors are shown, as well as for the subject factor which is nested within shape.

|  | Type | Shape | Subject(shape) |
| --- | --- | --- | --- |
| p-values | 0.0054 | $2.95 * 10^{-7}$ | 0.2674 |
| factor | fixed | fixed | random |

#### Source Phase-Shift Results

Here we present results of the time-delay analysis for the absolute joint-space representation and the task-space, and show that the time delays (*τ*_*ij*_) are not arbitrarily determined and expose correlations between the time delays, i.e. phase-shifts. Noteworthy results are highlighted. Essentially, the drawn shapes can be partitioned into 3D shapes (bent-ellipse or double bent ellipse) and planar shapes. An example figure showing the BE phase shifts is shown here, but see Appendix **?? B** for the results of the source phase-shifts for other shapes. The average results results are summarized in Table 3.

**Table 3:**
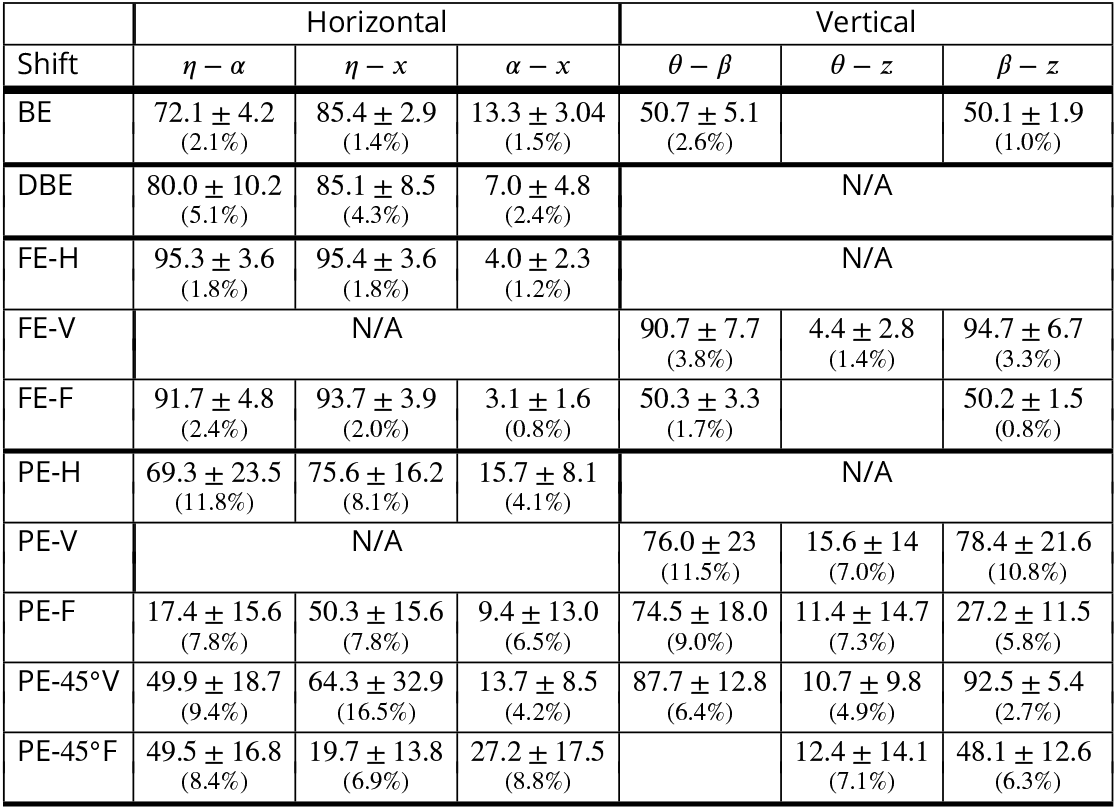
Average phase-shifts and SD (±) for the planar horizontal ellipse (PE-H), ellipse drawn in the sagittal plane (PE-V), ellipse drawn in the frontal plane (PE-F), ellipse drawn on a 45^°^ slanted plane with respect to the sagittal (PE-45^°^V) and ellipses drawn on a 45^°^ slanted plane with respect to the frontal plane (PE-45^°^F). The values in the parentheses stand for the SD as percentage of one cycle duration.

| Shift | Horizontal |  |  | Vertical |  |  |
| --- | --- | --- | --- | --- | --- | --- |
| | $\eta - \alpha$ | $\eta - x$ | $\alpha - x$ | $\theta - \beta$ | $\theta - z$ | $\beta - z$ |
| BE | 72.1 ± 4.2<br>(2.1%) | 85.4 ± 2.9<br>(1.4%) | 13.3 ± 3.04<br>(1.5%) | 50.7 ± 5.1<br>(2.6%) | | 50.1 ± 1.9<br>(1.0%) |
| DBE | 80.0 ± 10.2<br>(5.1%) | 85.1 ± 8.5<br>(4.3%) | 7.0 ± 4.8<br>(2.4%) | N/A | | |
| FE-H | 95.3 ± 3.6<br>(1.8%) | 95.4 ± 3.6<br>(1.8%) | 4.0 ± 2.3<br>(1.2%) | N/A | | |
| FE-V | N/A | | | 90.7 ± 7.7<br>(3.8%) | 4.4 ± 2.8<br>(1.4%) | 94.7 ± 6.7<br>(3.3%) |
| FE-F | 91.7 ± 4.8<br>(2.4%) | 93.7 ± 3.9<br>(2.0%) | 3.1 ± 1.6<br>(0.8%) | 50.3 ± 3.3<br>(1.7%) | | 50.2 ± 1.5<br>(0.8%) |
| PE-H | 69.3 ± 23.5<br>(11.8%) | 75.6 ± 16.2<br>(8.1%) | 15.7 ± 8.1<br>(4.1%) | N/A | | |
| PE-V | N/A | | | 76.0 ± 23<br>(11.5%) | 15.6 ± 14<br>(7.0%) | 78.4 ± 21.6<br>(10.8%) |
| PE-F | 17.4 ± 15.6<br>(7.8%) | 50.3 ± 15.6<br>(7.8%) | 9.4 ± 13.0<br>(6.5%) | 74.5 ± 18.0<br>(9.0%) | 11.4 ± 14.7<br>(7.3%) | 27.2 ± 11.5<br>(5.8%) |
| PE-45°V | 49.9 ± 18.7<br>(9.4%) | 64.3 ± 32.9<br>(16.5%) | 13.7 ± 8.5<br>(4.2%) | 87.7 ± 12.8<br>(6.4%) | 10.7 ± 9.8<br>(4.9%) | 92.5 ± 5.4<br>(2.7%) |
| PE-45°F | 49.5 ± 16.8<br>(8.4%) | 19.7 ± 13.8<br>(6.9%) | 27.2 ± 17.5<br>(8.8%) | | 12.4 ± 14.1<br>(7.1%) | 48.1 ± 12.6<br>(6.3%) |

#### 3D Shapes

We begin by observing the phase-shifts for the bent-ellipse shape (Figure 11). Interestingly, all subjects demonstrate similar behavior when inspecting certain phase-shifts. A constant phaseshift (72.08 ± 4.23 SD^2^) between the azimuth angles of the upper-arm and the forearm (*η* − *α*) appears in the first source (*s*_1_).

**Figure 9:**
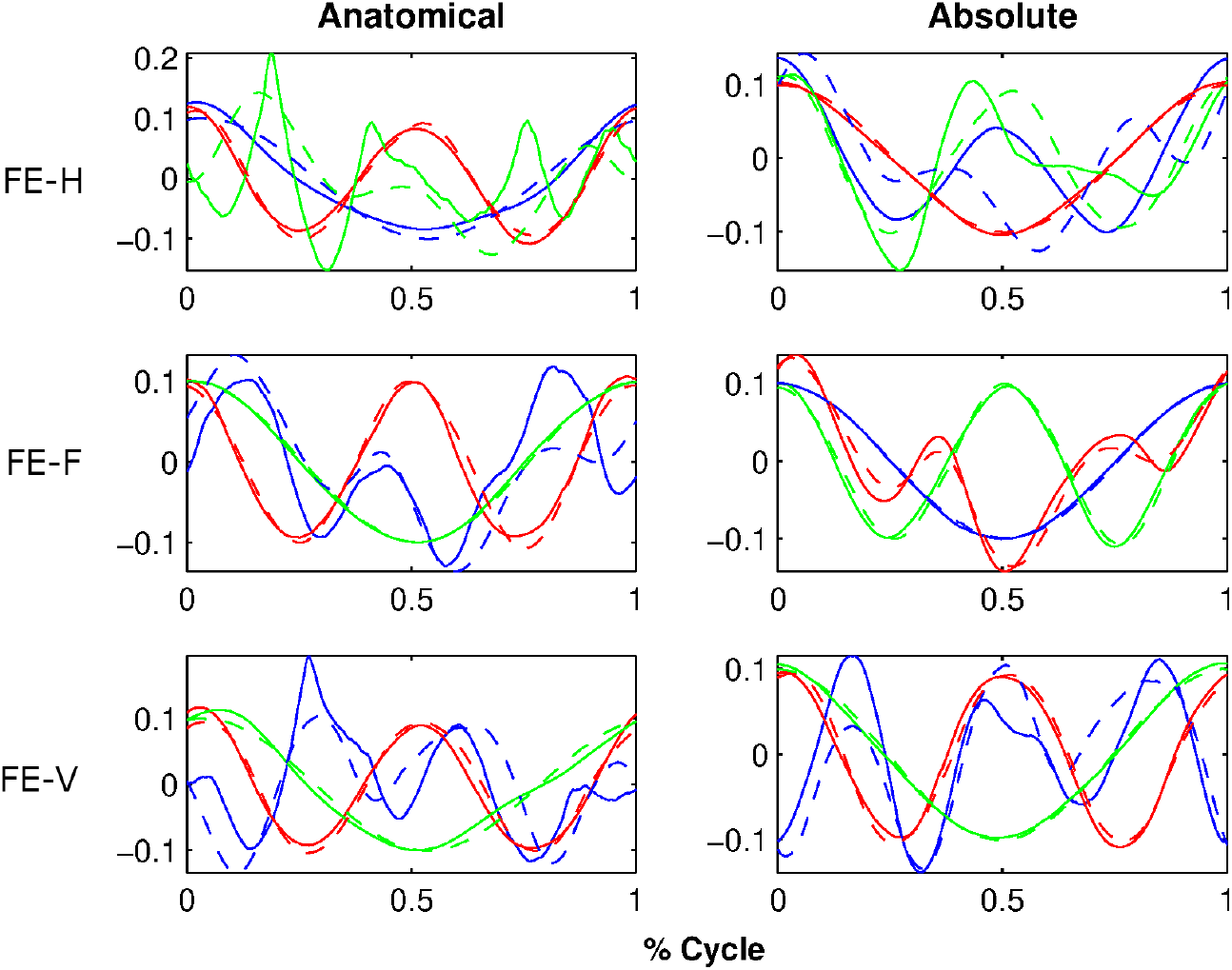
Extracted sources for the figure eight shapes at different orientation. FE-H is horizontal, FE-F is on the frontal plane, FE-V is on the sagittal plane. The rest of the details are as in Figure 8.

**Figure 10:**
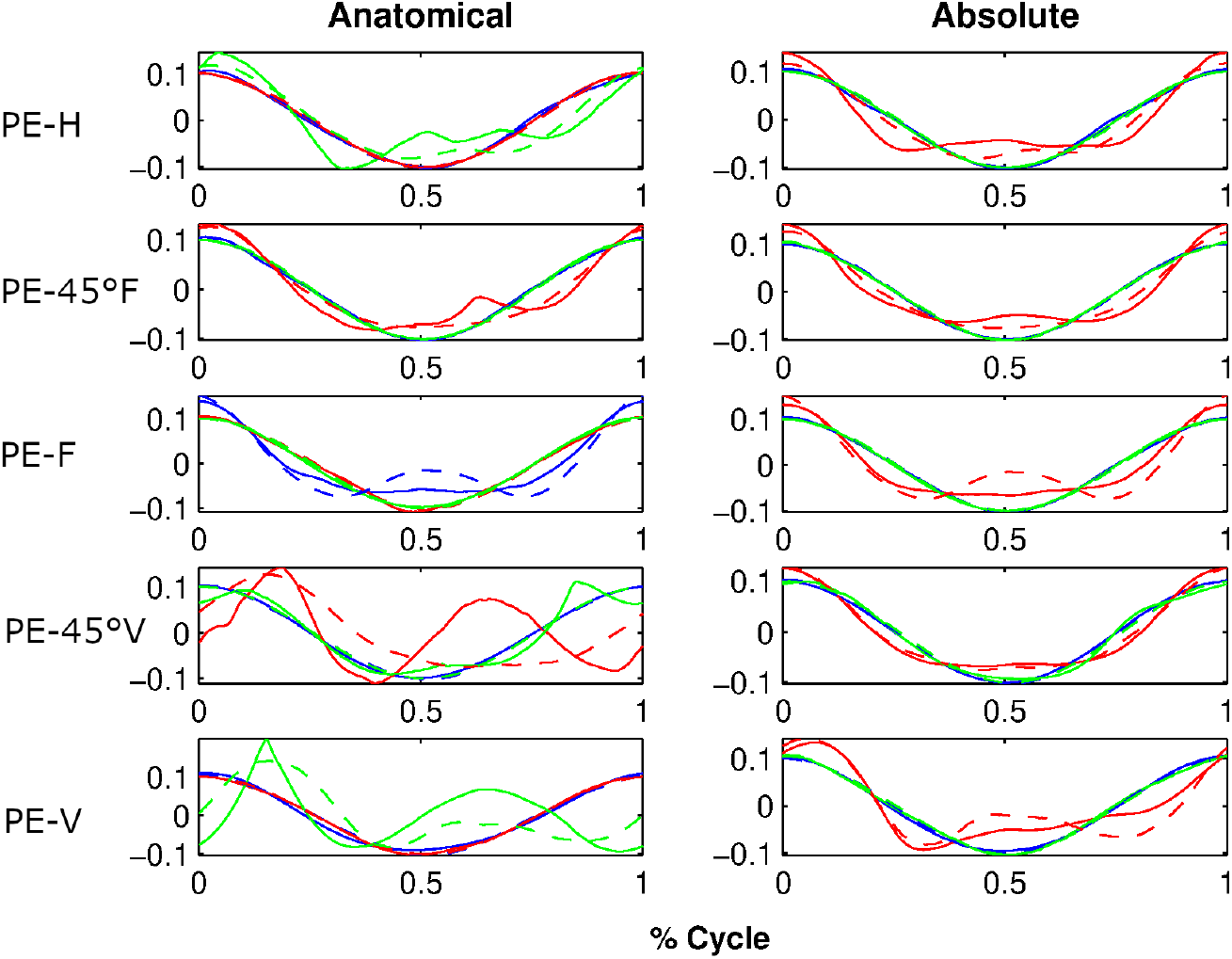
Extracted sources for the Planar Ellipses shapes at different orientations. PE-H is a horizontal ellipse, PE-45°F is tilted at 45 degrees with respect to frontal plane, PE-F is on the frontal plane, PE-45°V - 45 degrees with respect to sagittal, and PE-V is on the sagittal plane. The rest of the details are as in the Figure 8.

**Figure 11:**
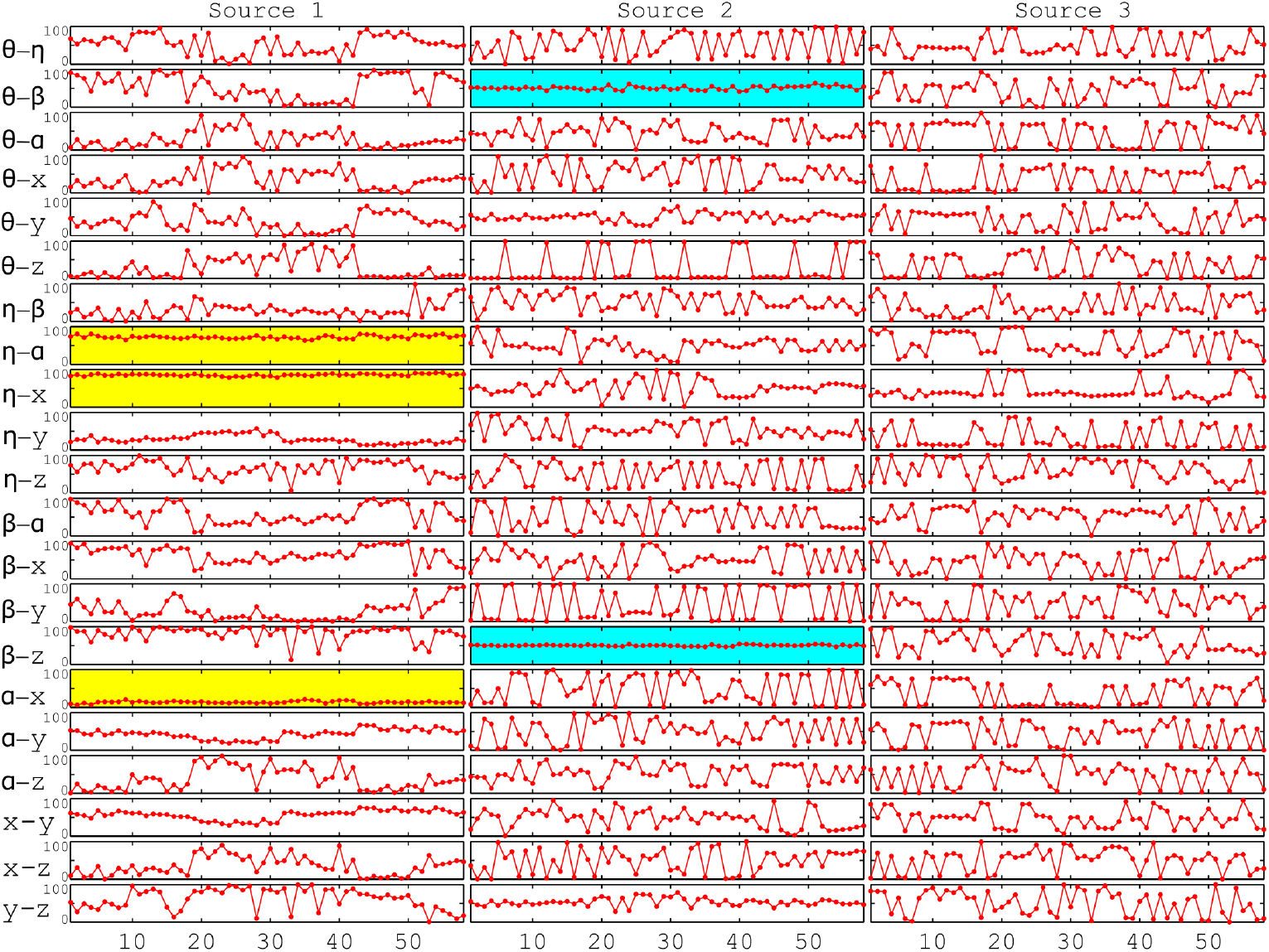
Bent Ellipse (BE) phase-shifts. The plot shows the phase shifts between the azimuth angle of the upper arm, *η*, azimuth angle of the forearm, *α*, elevation angle of the upper arm, *θ*, and elevation angle of the forearm *β*. Each source is represented by one column. For each source all possible phase-shifts between the DOFs are presented in rows. Each point in the plot represents one movement cycle. The horizontal axes denote the number of recorded cycles. Yellow and blue highlight behavior of interest, namely similar phase-shifts for all subjects. Yellow designates lateral related variables and blue highlights vertical related variables.

Similarly, the phase-shifts between the upper-arm’s azimuth angle and the lateral direction (*η*−*x*) and the forearm’s azimuth angle and the lateral direction (*α*−*x*), are constant for all subjects (highlighted with yellow in Fig 11 in Appendix **B**). Moreover, the second source (*s*_2_) exhibits phase locking between the upper-arm and forearm elevation angles (*θ* − *β*) and between the elevation angle of the forearm and the upwards direction (*β* − *z*) for all subjects (highlighted in blue in Fig 11).

The values of these phase-shifts are summarized in Table 3. It should be noted that these values can range between 0 and 200, since one normalized cycle for each shape contains 200 sample points. Thus, although the standard deviations in the table appear in units of sample points, it is better to view them as percentage of one cycle duration in order to assess how large they are. This means that standard deviations of 10 signify 5% of a cycle duration. That is, the standard deviations for the BE shape appearing in Table 3 are quite low and generally in the range of ∼ 3% and less.

Therefore, two main observations should be made. First, similar phase-shifts for all subjects are maintained not only within task or joint spaces, e.g., *η* − *α* but also across the spaces, e.g., *η* − *x*. Second, one source (*s*_1_) maintains the phase-shifts among the azimuthal related variables, whereas another source (*s*_2_) does the same for the elevational related variables. It is as if the variables of movement are distributed between two channels of control. One for azimuth and one for elevation. The third source does not seem to play a role in the azimuth or elevation control, but it may reflect the variations between the cycles and subjects.

This phenomenon repeats itself for the other shapes, however not always to its full effect as in the case of the bent-ellipse drawing. When we observe the results for the double bent ellipse drawing (Figure B.1), we see that the azimuth-related variables are phase locked, just like for the bent-ellipse shape. Again, the constant phase-shifts for the lateral (azimuth) related variables are handled by only one source (*s*_2_). However, the same is not obtained for the vertical related variables (also see Table 3).

#### Planar Shapes

When inspecting planar shapes, one can further divide these into shapes that contain movement only in one principal plane, (i.e. horizontal or sagittal) and shapes that are drawn on slanted planes which include movements in both the horizontal and vertical directions. The differences in the drawing planes observed in the type of phase-locks that emerge will be shown next. When drawing the figure eight shape in the horizontal plane (for context see Figure B.3 in Appendix **B**) one source (*s*_3_) maintains a phase lock among the azimuthal related variables.

This can be explained simply by the fact that there is no vertical component to the horizontal figure eight shape. Table 3 summarizes the phase shifts for the horizontal figure 8 in row FE-H.

Similarly, as would be expected, when drawing the figure eight in the sagittal plane, only vertical related variables are coupled (Figure B.4). These phase-shifts can be seen in Table 3 in row FE-V.

However, when the figure eight is drawn in the frontal plane, so that the shape has both lateral and vertical components, an identical phase locking pattern to that of the bent ellipse shape is observed (compare Figures 11 and C.3). The corresponding phase-shifts can be viewed in Table 3 in row FE-F. Once again, the SD in Table 3 should be considered as percentage of one period of a cycle. Thus, the SD for these phase-shifts are mostly less then ∼ 3% of a cycle period. Next, results for the planar ellipse are shown. In addition to the principal planes (horizontal, frontal and sagittal), ellipses were also drawn on 45^°^ slanted planes off the sagittal and frontal planes. The results for an example planar ellipse shape can be considered in Figure B.2 (in Appendix B). Although similar behavior as before can be detected, the variance values of the phase-shifts are higher with respect to the other shapes as can be observed in Table 3. That is, behavior during ellipse drawing was less consistent than for other shapes, with standard deviations of roughly 7.5% of one cycle period. It is not clear yet why results for the planar ellipse are worse than for the other cases. One possibility stems from the fact that the ellipse shape is not a con-straining shape. This fact may have led to larger variations in the arm configurations while the subjects were drawing this shape.

## Discussion

The question of how the nervous system controls and coordinates multi-joint movements is essential within neuroscience, as well as extremely important for developing rehabilitation robots and techniques that can ultimately improve function and movement abilities for individual subjects. In this work, we were concerned with the human arm kinematic redundancy problem. Specifically, we want to understand how the nervous system controls motion in joint-space to achieve a desired arm trajectory executed in task-space. This is far from a trivial question, since the redundancy of the DoFs available in the human arm relative to the 3D task-space DoFs renders the solution-space to be of very high dimensionality.

Thus, this work has two main results. First, we have shown that using a novel technique, we can identify sources (functional basis functions) that are shared by the task and joint spaces, when the joint space is represented in terms of an external reference frame (absolute angle representation). Second, we have shown that movement composition from basic sources and using the same sources at different levels of the motor hierarchy could possibly be the mechanism that the central nervous system uses to transform between task and joint spaces when solving the inverse kinematics of the arm while planning a particular movement. Furthermore, these results have shown that the set of sources, although delayed in time, are not arbitrarily delayed but are prescribed by couplings (or coordination) between the different degrees of freedom (DOFs). These results and future directions are discussed below.

### Dimensionality Reduction

The sources for the joint and task-spaces were strikingly similar when we applied the FADA source separation method. However, this similarity between the two source sets was very strong when the joint space was represented in an external frame of reference (or absolute angles/coordinates), but significantly less strong when the joint-space was represented in anatomical/relative coordinates. It should be noted that this dimensionality reduction method is *unsupervised* and no assumptions were made with respect to the functional form, *ϕ*, of the sources.

The fact that we were able to represent the joint-space with a similar set and number of sources as those of the task-space (three in both cases) means not only that the intrinsic dimensionality of the joint-space is the same, but that these spaces share some geometrical properties and structure. Since these two different spaces share a single set of sources, we were able to show that one can go back and forth between the joint and task spaces. The sources themselves can serve as “mediators” (channels) between the configuration and task spaces. This phenomenon may be used by the CNS as a mechanism to solve the inverse kinematics problem.

It should be noted that the suggested solution is general, in the sense that it does not require the underlying sources to have specific properties. The only requirement is that the joint and task spaces share the same set of sources. As long as this requirement is satisfied, they can be integrated into the inverse kinematics solution regardless of what those sources look like or what method is used to compute the sources. Thus, the solution is independent of the specific methodology used for acquiring them. However, this work only lays the groundwork for the solution and is not yet complete. The correct time-delays for the sources in joint-space need to be identified. We know that these time-delays are not arbitrary and are constrained by the specific shapes that are drawn and by the timing imposed on the sources by the task-space.

### Phase-Shifts & Coordination

The individual time delays of each source in joint-space need to be determined to produce the correct trajectories in that space. We have shown that different DOFs both in the joint-space and across the joint and task spaces are *coordinated* with specific time-delays. For a specific pairing of variables, we showed that specific phase-shifts exist for all subjects and trials.

Specifically, we have shown that variables related to lateral movement (azimuth) tend to be phase-locked by one source, and variables related to the elevation (i.e. vertical) are coordinated by another source. Indeed, this result comes from a completely computational process. However, this may reveal elements of the way the solution is organized. It is possible that coordination isn’t constrained to a specific configuration space, and variables in the task and joint spaces could potentially be coupled. This notion could lead the way to a solution that uses time-delays in task-space to determine delays in joint-space. Note that this is a direct consequence of the specific selected reference frame, as will be discussed in the next section.

### Reference Frames

Previously, numerous studies (***Soechting and Ross, 1984***; ***Soechting, 1982***; ***Worringham and Stelmach, 1985***; ***Worringham et al., 1987***) showed a high preference for representing arm configurations in an extrinsic, or absolute, reference frame. This preference fits well with the results from our analyses, where the absolute joint-space sources are a good fit for the task-space sources.

***Pozzo et al. (1998***) have argued that given that gravity is a force that is sensed by both the vestibular and proprioceptive systems, therefore guidance of a body segment in space requires the CNS to use an absolute reference frame within which the external positions and displacements of the whole body could be estimated. In fact, there is evidence that the nervous system does have an internal model of gravity that can inform movements (***McIntyre et al., 2001***).

Similarly, in an extensive review in which a variety of different tasks were considered, ***Soechting and Flanders (1992***) noted that in all coordinate systems one axis was defined by the gravitational vertical, and another was defined by a sagittal horizontal axis. This may be due to the neural control stategy whereby human movements in a three-dimensional world are dominated by the force of gravity and by the effect of visual horizon on motion planning. These authors therefore concluded that this common, spatial frame of reference might aid in the exchange of information between brain regions. Indeed, electrophysiological data obtained from neural recordings in the superior colliculus and motor cortex have suggested that this is the case, with neural activity in both structures appearing to encode movement kinematics, specifically movement direction.

Meanwhile, there is work in monkey models showing reach planning is performed in eye-coordinates rather than arm-coordinates, and this is essential to hand-eye coordination (***Batista et al., 1999***). Additionally, there is some evidence that at least in some brain regions, reaching tasks are represented in body-coordinate frames (***Lacquaniti et al., 1995***) which seems contrary to our results. However, the literature is quite mixed, with some researchers reporting results that seem to imply a neural preference for representing motion in hand or body-coordinates (***Soechting et al., 1990a***; ***Wang and Spelke, 2000***; ***McIntyre et al., 1998***), and others reporting preference for eye or world-coordinates (***Bosco et al., 2000***; ***Poljac and Van Den Berg, 2003***), and others reporting multiple coordinate systems may be used for different representations or in different brain regions (***Carrozzo et al., 2002***; ***Graziano et al., 1994***). Perhaps, as was proposed in ***Andersen et al. (2007***) there is a shared coordinate system used, with differentiated gains used to modulate or transform between coordinates in eye or body space. Our results seem to support the preference for world-coordinates at least in unconstrained arm movements, but this is subject to further investigation.

Importantly, our results support previous work, where ***Soechting and Ross (1984***) and ***Flanders and Soechting (1990***) showed that arm movements seem to be represented by yaw and elevation angles. Specifically, it was suggested by ***Flanders and Soechting (1990***) that target distance and elevation are used to compute arm elevation, and that target azimuth is used to compute arm yaw. Note the similarity between their results to the results we obtained, where one source controls the vertical variables and a second source controls the horizontal variables. At present we do not think that the third source has a specific role, and it is postulated that it accounts for the movement variability between different subjects and cycles.

### Compositionality

There is also a growing body of literature that supports the idea of compositionality (***Giszter, 2015***; ***d’Avella and Bizzi, 2005***; ***Flash and Hochner, 2005***). Compositionality is a concept, by which rather than controlling individual muscles, etc. the nervous system “composes” complex movements from underlying behaviors or control units, known as motor primitives or synergies, where multiple muscles, nerves etc. are coupled in a specific way. These primitives are specifically parameterized or optimized to fit motor needs for specific tasks, such as locomotion (***Ivanenko et al., 2004***; ***Merkle et al., 1998***) or arm movements (***d’Avella et al., 2006***; ***d’Avella and Lacquaniti, 2013***; ***Flash and Berthoz, 2021***). Though synergies are somewhat debated, there is evidence for muscle synergies (***Bizzi et al., 1995***; ***Tresch et al., 1999***; ***Mussa-Ivaldi et al., 1994***; ***Ivanenko et al., 2004***). In a sense, the sources we’ve shown here could be considered spatio-temporal synergies (or neural control synergies/motor primitives) and help provide evidence for synergies of movement coordination. However, the question remains whether the muscle synergies are wholly a product of the neural resolution of redundancy using underlying sources or based on biomechanical or spinal constraints.

Previous work has shown that already at birth or just shortly after the spinal and supraspinal networks seem to have intact motor primitives (***Yang et al., 2019***; ***Dominici et al., 2011***). Our results suggest that these motion primitives are represented neurally by means of basis functions or sources, where joint movements are coordinated using a representation of sources (***Yang et al., 2019***). However, tuning of this network is not complete at birth, and the coefficients for producing movements in either the task or joint spaces are *learned* during maturation until network convergence. Similar ideas appeared for the two-thirds power law, where it was observed that the law substrates exist quite early in development but the movement trajectories don’t converge toward exhibiting their mature characteristics until the age of approximately 12 years old (***Viviani and Schneider, 1991***). Additionally, the same suggestion was made for the law of intersegmental coordination during locomotion by ***Bleyenheuft and Detrembleur (2012***). In both cases, it was suggested that the network exists at a very early age, but is tuned over time, as described in the work by ***Dominici et al. (2011***) where this was evaluated with newborn babies. Future work should further explore such tuning.

Finally, this work provides further evidence that the CNS exploits sources (or basis functions) and an absolute reference frame to devise a strategy for solving the inverse kinematics problem. Ultimately, the shape of the specific sources used by the nervous system is still a subject of research. The FADA-based sources we computed here may be close to those represented in the CNS, or they may be closer to pure sinusoids, as used by ***Barliya et al. (2009***) in their oscillator model of the intersegmental coordination. Future work will continue to focus on characterizing the sources and the time shifts between them more carefully, as well as considering their representations within the nervous system.

## Conclusion

In this work we have presented theoretical work based on our kinematic analysis of experimentally recorded arm movements, showing that a single set of sources could be used to represent movement trajectories in both the task and joint-spaces. In particular we used this observation to mathematically formulate a solution to the inverse kinematics problem. We showed that a set of three constituent sources could be extracted from movement data and, depending on the reference frame, could be used to represent both joint and task-space movements. Importantly, our analysis showed that using an external, absolute angle representation yielded significantly closer correlation between the joint and task-space sources, and therefore better transferabil-ity. Additionally, we investigated the time-delays between degrees of freedom, and found that there are specific couplings that exist and can be identified through our source decomposition approach. We also found essentially one source related to horizontal (azimuth) variables and one source related to vertical (elevation) variables. Finally, we believe that this work has implications for our understanding of the operation and coordination of the central nervous system in the planning of movement, and therefore continuing research towards this end is important for our understanding of neural mechanisms of human movement generation and motor-disorders and pathologies.

## Acknowledgments

Nili Krausz was supported by a Zuckerman STEM Leadership Postdoctoral Fellowship. Tamar Flash and Martin Giese were supported, in part, by the CRCNS US-Israeli-German Collaborative Research grant in computational neuroscience. Tamar Flash was also supported by grants from the Estate of Naomi K. Shapiro, the Rolf and Alice Wiklund Parkinson’s Disease Research Fund, the Steven Gordon and Levine Foundations, and the Rudolph and Hilda U. Forchheimer Foundation. Martin A Giese was also funded by ERC 2019-SyG-RELEVANCE-85649.

## Footnotes

1 A linear transformation *L* between normed vector spaces *X* and *Y* for which there exists some *M* > 0 such that for all *v* ∈ X, ‖*Lv*‖_*Y*_ ≤ *M*‖*v*‖_*X*_

2 Hereafter, all average values will be accompanied with standard deviation (SD) values, unless mentioned otherwise and the sign (SD) will be omitted.

## Notes

### Competing Interest Statement

The authors have declared no competing interest.

### Summary of Updates

We have slightly edited for improved clarity

