## Appendices for "Human Arm Redundancy - A New Approach for the Inverse Kinematics Problem"

#### Appendix A) Inverse of $A(\xi)$ matrix

Since $A(\xi)$ is a $3\times 3$ matrix, its inverse is easily computed. The inverse of a general $3\times 3$ matrix $M$ is given by

$$M^{-1} = \frac{1}{|M|}\begin{pmatrix} \begin{vmatrix} m_{22} & m_{23} \\ m_{32} & m_{33} \end{vmatrix} & \begin{vmatrix} m_{13} & m_{12} \\ m_{33} & m_{32} \end{vmatrix} & \begin{vmatrix} m_{12} & m_{13} \\ m_{22} & m_{23} \end{vmatrix} \\ & & \\ \begin{vmatrix} m_{23} & m_{21} \\ m_{33} & m_{31} \end{vmatrix} & \begin{vmatrix} m_{11} & m_{13} \\ m_{31} & m_{33} \end{vmatrix} & \begin{vmatrix} m_{13} & m_{11} \\ m_{23} & m_{21} \end{vmatrix} \\ & & \\ \begin{vmatrix} m_{21} & m_{22} \\ m_{31} & m_{32} \end{vmatrix} & \begin{vmatrix} m_{12} & m_{11} \\ m_{32} & m_{31} \end{vmatrix} & \begin{vmatrix} m_{11} & m_{12} \\ m_{21} & m_{22} \end{vmatrix} \end{pmatrix}$$

where $|\cdot|$ denotes the determinant of a matrix. Thus, $A^{-1}(\xi)$ is of the form

$$A^{-1}(\xi) = \frac{1}{|A|}\begin{pmatrix} m_{11}(\xi) & m_{12}(\xi) & m_{13}(\xi) \\ m_{21}(\xi) & m_{22}(\xi) & m_{23}(\xi) \\ m_{31}(\xi) & m_{32}(\xi) & m_{33}(\xi) \end{pmatrix}$$

and the elements $m_{ij}$ after some simple algebraic manipulations are given by,

$m_{11} = a_{22}a_{33}e^{-2\pi i\xi(\tau_{22}+\tau_{33})} - a_{23}a_{32}e^{-2\pi i\xi(\tau_{23}+\tau_{32})}$
$m_{12} = a_{13}a_{32}e^{-2\pi i\xi(\tau_{13}+\tau_{32})} - a_{12}a_{33}e^{-2\pi i\xi(\tau_{12}+\tau_{33})}$
$m_{13} = a_{12}a_{23}e^{-2\pi i\xi(\tau_{12}+\tau_{23})} - a_{13}a_{22}e^{-2\pi i\xi(\tau_{13}+\tau_{22})}$
$m_{21} = a_{23}a_{31}e^{-2\pi i\xi(\tau_{23}+\tau_{31})} - a_{21}a_{33}e^{-2\pi i\xi(\tau_{21}+\tau_{33})}$
$m_{22} = a_{11}a_{33}e^{-2\pi i\xi(\tau_{11}+\tau_{33})} - a_{13}a_{31}e^{-2\pi i\xi(\tau_{13}+\tau_{31})}$
$m_{23} = a_{13}a_{21}e^{-2\pi i\xi(\tau_{13}+\tau_{21})} - a_{11}a_{23}e^{-2\pi i\xi(\tau_{11}+\tau_{23})}$
$m_{31} = a_{21}a_{32}e^{-2\pi i\xi(\tau_{21}+\tau_{32})} - a_{22}a_{31}e^{-2\pi i\xi(\tau_{22}+\tau_{31})}$
$m_{32} = a_{12}a_{31}e^{-2\pi i\xi(\tau_{12}+\tau_{31})} - a_{11}a_{32}e^{-2\pi i\xi(\tau_{11}+\tau_{32})}$
$m_{33} = a_{11}a_{22}e^{-2\pi i\xi(\tau_{11}+\tau_{22})} - a_{12}a_{21}e^{-2\pi i\xi(\tau_{12}+\tau_{21})}$

The determinant of $A$ is

$$\begin{aligned} |A| \quad = \quad & -a_{13}a_{22}a_{31}e^{-2\pi i\xi(\tau_{13}+\tau_{22}+\tau_{31})} + a_{12}a_{23}a_{31}e^{-2\pi i\xi(\tau_{12}+\tau_{23}+\tau_{31})} \\ + \quad & a_{13}a_{21}a_{32}e^{-2\pi i\xi(\tau_{13}+\tau_{21}+\tau_{32})} - a_{11}a_{23}a_{32}e^{-2\pi i\xi(\tau_{11}+\tau_{23}+\tau_{32})} \\ - \quad & a_{12}a_{21}a_{33}e^{-2\pi i\xi(\tau_{12}+\tau_{21}+\tau_{33})} + a_{11}a_{22}a_{33}e^{-2\pi i\xi(\tau_{11}+\tau_{22}+\tau_{33})} \end{aligned}$$

Finally, $A$ is invertible if and only if $|A| \neq 0$.

### Appendix B) Phase Shift Plots

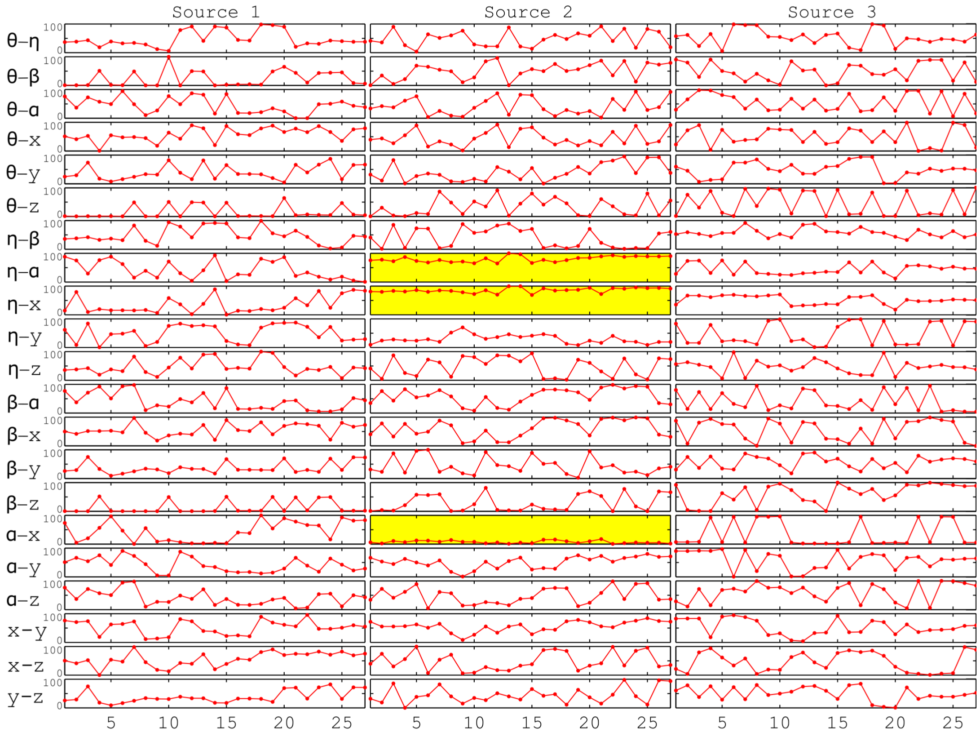


Figure B.1: Double Bent Ellipse - Phase-shifts. See Figure 11 for details.

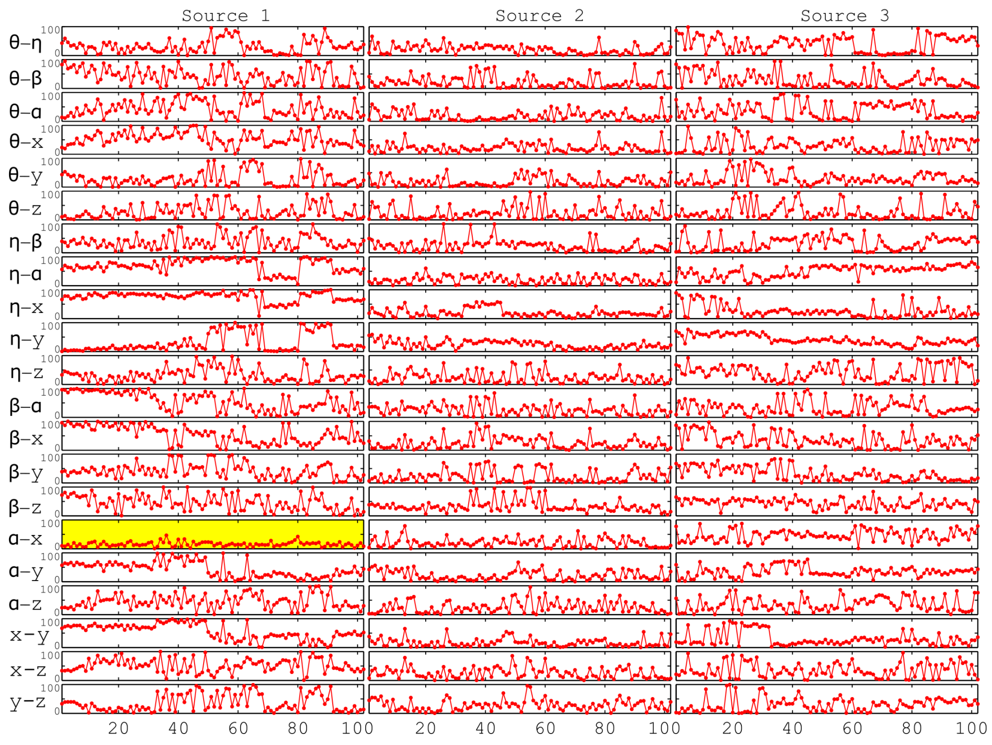


Figure B.2: Planar Ellipse - phase-shifts. See Figure 11 for details

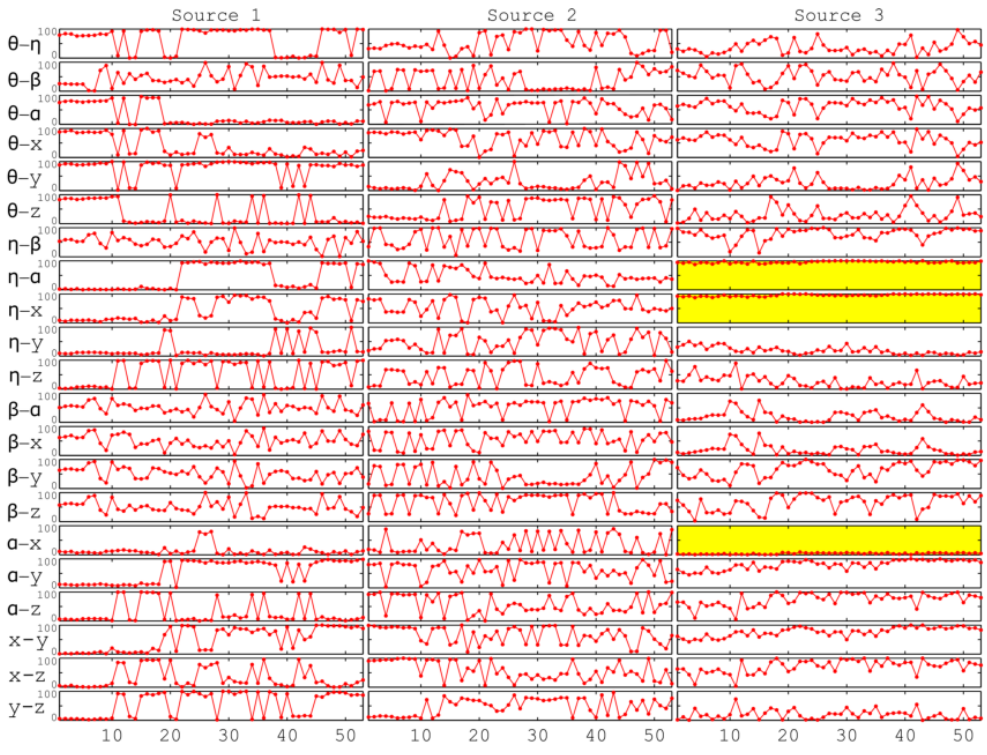


Figure B.3: Figure Eight - phase-shifts. See Figure 11 for details

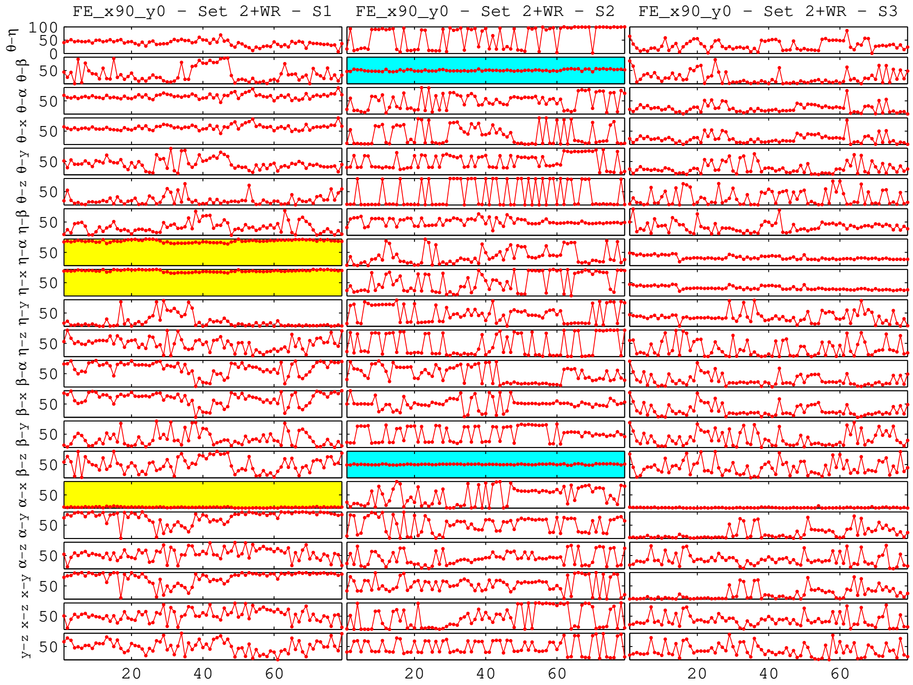


Figure B.4: Figure 8, rotated $90°$ about $x$-axis - Phase-shifts. See Figure 11 for details.

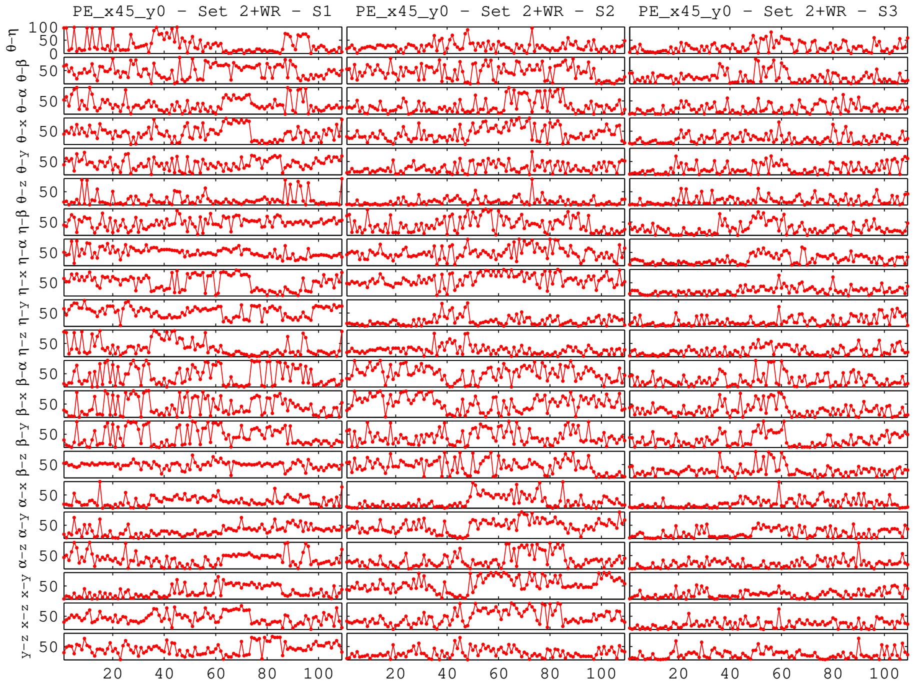


Figure B.5: Ellipse rotated $45°$ about $x$-axis - Phase-shifts. See Figure 11 for details.

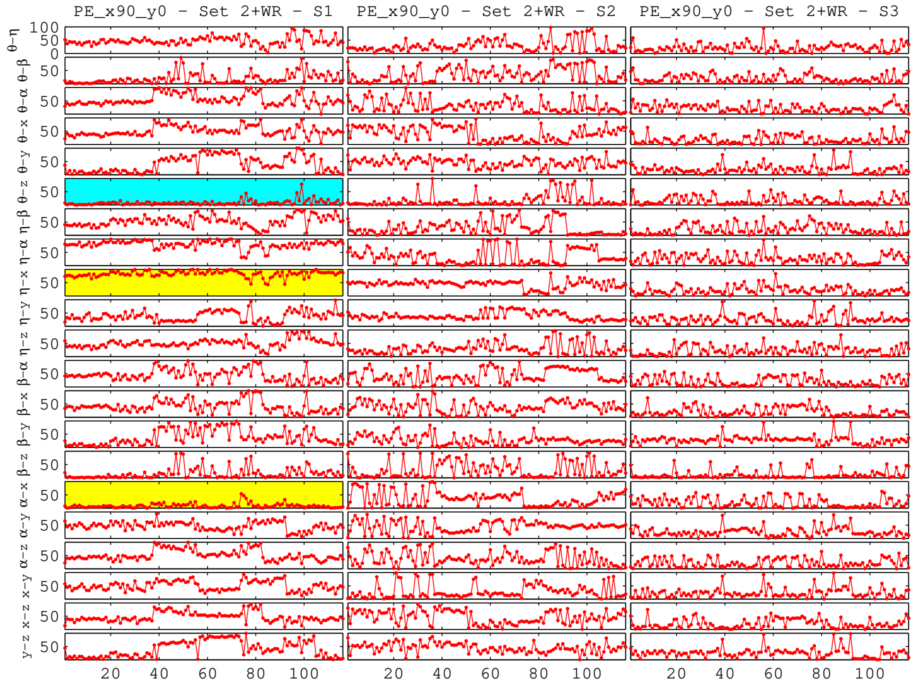


Figure B.6: Ellipse rotated $90°$ about $x$-axis (frontal plane) - Phase-shifts. See Figure 11 for details.

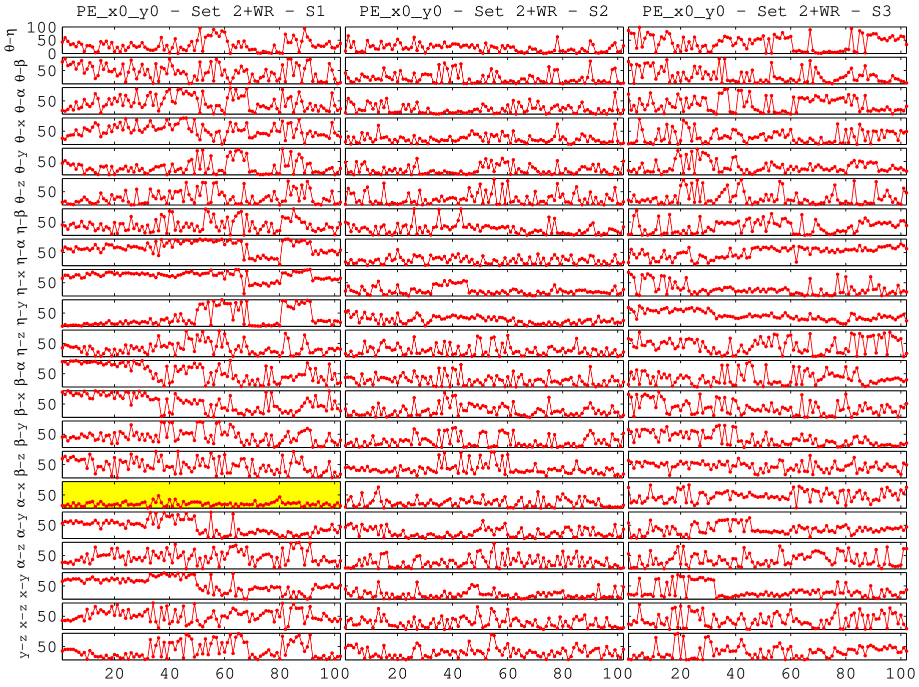


Figure B.7: Horizontal Ellipse - Phase-shifts. See Figure 11 for details.

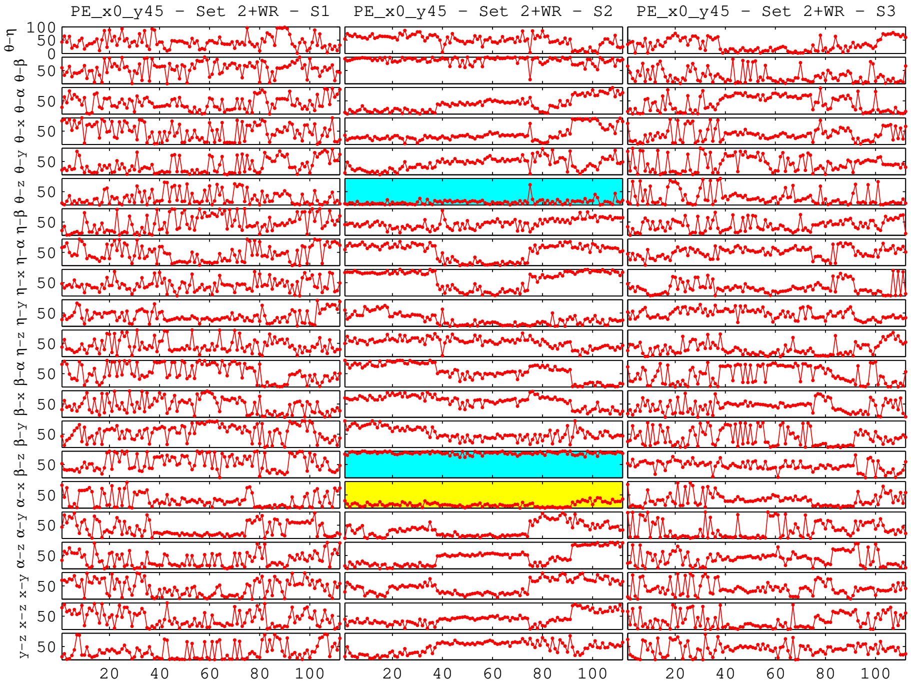


Figure B.8: Ellipse rotated 45° about $y$-axis - Phase-shifts. See Figure 11 for details.

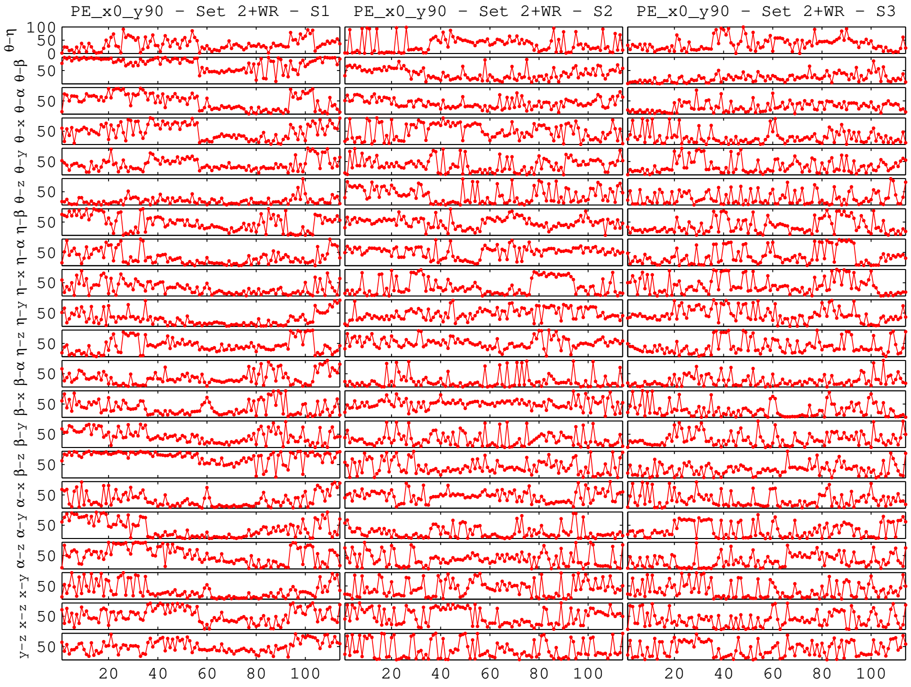


Figure B.9: Ellipse rotated 90° about $y$-axis (sagittal plane) - Phase-shifts. See Figure 11 for details.
